# Biologically Driven *In Vivo* Occlusion Design Provides a Reliable Experimental Glaucoma Mouse Model

**DOI:** 10.1101/2024.01.18.576306

**Authors:** Eunji Hong, Qinglong Wang, Ruyi Yang, Feng Tian, Ryan Donahue, Xianyang Li, Christopher Glynn, Sophia Tsekov, Chen Lin, Mengwen Xiong, Sizhe Huang, Songlin Zhou, Lan Yao, Jifu Tan, Ying Wang, Zhigang He, Siyuan Rao, Qianbin Wang

**Affiliations:** Department of Biomedical Engineering, Binghamton University, State University of New York, Binghamton, NY, 13902, USA; Department of Neurology, Beth Israel Deaconess Medical Center, Harvard Medical School, Boston, MA, 02115, USA; F.M. Kirby Neurobiology Center, Boston Children’s Hospital, Harvard Medical School, Boston, MA, 02115, USA; Department of Biomedical Engineering, University of Massachusetts, Amherst, MA, 01003, USA; Small Scale System Integration and Packaging (S3IP), Binghamton University, State University of New York, Binghamton, NY, 13902, USA; Department of Mechanical Engineering, Binghamton University, State University of New York, Binghamton, NY, 13902, USA; Integrative Neuroscience Program, State University of New York at Binghamton, Binghamton, NY, 13902, USA

**Keywords:** Viscobeads, Glaucoma, Intraocular Pressure, Neurodegeneration

## Abstract

Fluid transport through the trabecular meshwork is a major regulator of intraocular pressure in healthy and glaucomatous eyes. Existing microbead occlusion models enable *in vivo* study of pressure-related glaucomatous pathology, but their long-term reliability is limited by inadequate bead-tissue interface design. Inspired by the graded porous architecture and fluid-transport function of the trabecular meshwork, we develop an injectable Viscobeads platform for sustained modulation of aqueous humor outflow. These composite microbeads integrate a non-degradable polystyrene core for structural support with a biodegradable poly(lactic-co-glycolic acid) viscoelastic shell that improves mechanical adaptation to heterogeneous trabecular meshwork fenestrations. This biologically informed design enhances outflow obstruction and enables stable IOP elevation for at least 8 weeks after a single injection. In mice, Viscobeads induce sustained ocular hypertension (average 21.4 ± 1.39 mm Hg) and lead to a 34% loss of retinal ganglion cells by day 56. Pattern electroretinogram and flash visual evoked potential measurements further reveal early retinal ganglion cell dysfunction followed by later visual pathway impairment. Beyond disease induction, this platform supports *in vivo* gene screening for retinal ganglion cell survival and identifies altered sleep behavior during glaucoma progression. This work establishes a materials-driven strategy for chronic ocular hypertension modeling and therapeutic evaluation.

## 1 Introduction

Fluid flow transport through biological tissues plays important roles in regulating pressure ^[1]^, morphogenesis ^[2]^, homeostasis and pathogenesis ^[3]^. For instance, the intraocular pressure (IOP) is regulated by aqueous humor outflow through the trabecular meshwork (TM), comprised of variably sized intertrabecular fenestrations ^[4]^. Occlusion of the trabecular meshwork impedes the outflow drainage and consequently leads to IOP elevation and glaucomatous neurodegeneration, which is a progressive and irreversible neuropathic condition in the visual system ^[5]^.

Glaucoma therapeutic development requires a reliable *in vivo* model to capture the pathophysiology associated with chronic elevation of IOP. Laboratory mouse models provide a useful *in vivo* system to study the pathophysiology of human diseases with various genetic approaches; however, current transgenic mouse models, such as DBA/2J, are limited by inconsistent IOP elevation, which leads to large variations in neural mechanism investigation. Among experimental glaucoma animal models, engineered occlusion models with injectable microscale biomaterials to manipulate the aqueous humor (AH) outflow have attracted attention due to their relatively straightforward operations and rapid IOP elevation ^[6]^. Polystyrene (PS) or latex microspheres with either uniform or mixed sizes have been translated into commercial products for research communities. However, these models either require multiple injections due to poor retention of the injected microbead inside the tissue or result in variable IOP elevation across different studies, ranging from 13 to over 30 mm Hg^[7]^. We hypothesize that these limitations primarily stem from the inadequate design of microbeads to accommodate the complex TM architecture and optimize material-tissue interactions. Alternatively, the magnetic microbead method offers a different approach by using an external magnet to control microbead distribution within the anterior chamber, potentially improving retention and IOP consistency. While promising, this method introduces additional procedural complexity and requires specialized equipment, which may limit its widespread adoption in research settings.

Biophysical consideration of the material-tissue interface within the TM architecture imparts a basis to design microbead-based approaches to modulate AH outflow and recapitulate elevated IOP and retinal degeneration. The microbeads occlusion model design consists of two main considerations related to stable IOP elevation: the effective blockage of AH in the trabecular meshwork, which causes acute and effective IOP elevation, and the retention of injectable beads in TM tissues over the long term to prevent a return to normal IOP ^[8]^. Analysis revealed that the efficacy of occlusion is determined by infiltration of the injectable fillers across the TM architecture, a gradient of diverse pore sizes, while the long-term IOP elevation stability is affected by the interfacial interaction of beads and tissues ^[9]^.

Under dynamic fluidic transport conditions *in vivo*, rigid materials provide mechanical support to prevent degradation and structural collapses over time, and viscous surfaces provide strong interactions with tissues to retain beads against shear forces and turbulence in AH flow ^[10]^. Current PS and latex microsphere models, which only consist of rigid materials without viscous surfaces, exhibit limited retention time and consequently require multiple injections to maintain IOP elevation over time ^[7a, 11]^. Alternatively, the two-step injections of viscous materials like hyaluronic acid (HA) followed by polymeric microbeads, can only achieve transient IOP elevation due to the evacuation of the viscous materials ^[12]^. Insufficient materials interfacial design introduces undesirable variance *in vivo* and undermines the study of complex pathological mechanisms over chronic assessment.

Here, we present a “Viscobeads” technology, which regulates fluidic transport to reconstitute glaucomatous IOP elevation *in vivo*. To address the mismatches of materials-tissue interfaces, we synthesized a set of viscoelastic polymeric beads consisting of a rigid PS skeleton to provide mechanical support and a biodegradable poly(lactic- co-glycolic acid) (PLGA) soft surface, termed PS/PLGA Viscobeads. After a single injection, variously sized Viscobeads immediately block the graded sizes of the porous structures, sustained by the viscoelastic interface, to induce and maintain IOP elevation. By optimizing the size distribution of the Viscobeads to match the TM porous architecture, we observed that high IOP (≥ 15 mm Hg persisting over three consecutive days) elevation was sustained for over 50 days. Corresponding to the prolonged high IOP maintenance window, we observed over 34% retinal ganglion cells (RGC) loss at 56 days post-injection through cellular morphology assessment. Additionally, changes in RGC function and the integrity of the visual pathway during glaucoma progression were detected using pattern electroretinogram (pERG) and flash visual evoked potential (VEP). Since retinal degeneration represents an important feature of experimental glaucoma, we tested a series of gene candidates in our Viscobeads model and identified protective genes for further prevention approaches for glaucomatous neurodegeneration. Our early version of this design has demonstrated inducible and consistent IOP elevation in mouse models ^[13]^. It proves to be an effective approach for establishing an experimental glaucoma model and studying the genetic manipulation of identified transcription factors in RGC protection.

## 2 Results

### 2.1 Design of Viscobeads Matching the Sizes of Trabecular Meshwork Fenestrations

In the conventional AH drainage pathway, AH is secreted by the ciliary body in the posterior chamber, circulates within the anterior chamber, and exits the eye through the TM ^[4a]^ (**Fig. 1A-B**). In cases of open-angle glaucoma, TM obstruction impairs AH outflow, leading to increased IOP (**Fig. 1C**). To mimic this natural pathological process, we injected Viscobeads to modulate fluid transport and reconstitute glaucomatous IOP elevation in vivo. We first synthesized PS/PLGA Viscobeads with a controllable size distribution using the emulsion-based method ^[14]^ (**Supplementary methods 1 and 2**). These beads feature a stiff PS core and a viscoelastic PLGA surface, attributes crucial for prolonging retention in the TM. Hydrophobic PS functions as a rigid skeletal structure to ensure long-term stability, while hydrophilic and biodegradable PLGA provides viscoelastic surfaces after degradation *in vivo*. Upon intracameral injection of Viscobeads into anesthetized mice with a bubble tamponade in the anterior chamber (**Supplementary method 3**), the denser microbeads rapidly accumulated at the iridocorneal angle due to the liquid-air interface (**Fig. 1C, Fig. S1**). They were effectively retained across the structurally graded TM pores, obstructing aqueous humor drainage. To track the trajectory of Viscobeads post-injection, we incorporated Rhodamine B into the PLGA oil phase during their preparation (**Supplementary method 2**). Fluorescence microscopy confirmed effective blockage at the iridocorneal angle in the mouse anterior chamber of the eye (**Fig. 1D**).

**Fig. 1:**
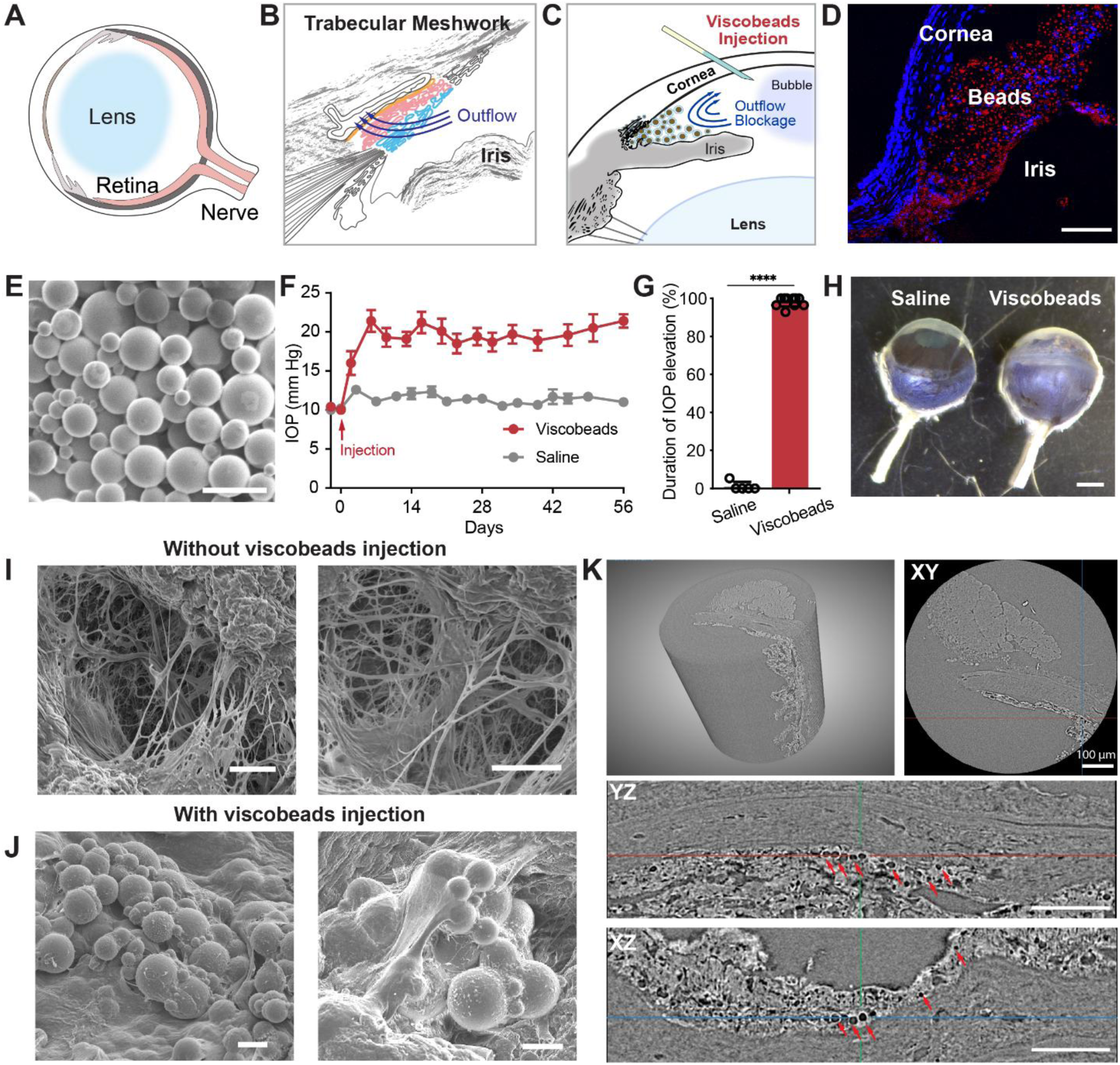
Bioinspired Design Injectable Viscobeads with Heterogeneous Size Distribution and Biodegradable Surface Properties. (A) Illustrations of the mouse eye, with the anterior segment comprising the cornea, iris, ciliary body, and lens. The ciliary body secretes aqueous humor, which circulates (blue arrows) within the anterior chamber before draining from the eye. (B) Enlarged view of the aqueous humor outflow pathway (blue arrows) in the iridocorneal angle. In the conventional pathway, the aqueous humor traverses the trabecular meshwork, first passing through the uveal meshwork (UM, highlighted in blue), then the corneoscleral meshwork (CM, highlighted in pink), and finally, the juxtacanalicular tissue (JCT, highlighted in yellow) before entering Schlemm’s canal. (C) An illustrative depiction of the injectable PS/PLGA Viscobeads technique. Viscobeads were injected intracamerally into the iridocorneal angle of the eye, blocking the aqueous humor outflow pathway. (D) A representative fluorescence image revealing the accumulation of rhodamine B-labeled PS/PLGAViscobeads at the mouse iridocorneal angle. Scale bar: 100 μm. (E) A representative SEM image of PS/PLGA Viscobeads displaying a size distribution ranging from 1 to 20 µm. Scale bar: 20 µm. (F) Intraocular pressure (IOP) in mice before and after the injection of PS/PLGA Viscobeads or the saline group. Data are shown as mean ± S.E.M. (G) The duration of high IOP (≥ 15 mm Hg) in mice injected with PS/PLGA Viscobeads and the saline group. Data are shown as mean ± SD. n=5 eyes for saline; 10 eyes for PS/PLGA Viscobeads group. Statistical significance used an unpaired two- tailed t-test. t(13) = 71.34, ****p<0.0001. (H) Representative photograph of eyes extracted 8 weeks post-injection of Viscobeads (glaucoma) and saline. (I) Representative scanning electron microscopy (SEM) images showing the trabecular meshwork’s gradient porous structure, with pore sizes ranging from approximately 1–20 μm. Scale bar: 10 μm. (J) Representative SEM images illustrating the entrapment of larger Viscobeads on the surface of the initial interlaminar spaces of the trabecular meshwork. (K) Representative 3D reconstruction of Micro-CT imaging of gel- embedded TM samples, shown in XY, YZ, and XZ views. Darker dots (red arrows) indicate the entrapment of Viscobeads within the tissue. Scale bar: 100 μm.

By adjusting the power, duty cycle, and ultrasound duration during oil-in-water synthesis^[15]^, we produced heterogeneous Viscobeads ranging from 1 to 20 μm in size (**Fig. 1E**). We then performed tonometry on both the Viscobeads and saline injection groups to assess IOP elevation. As Viscobeads rapidly obstructed the trabecular meshwork’s porous structure, IOP increased sharply to 21.4 ± 1.39 mm Hg by day 7 post-injection and remained elevated at approximately 20 mm Hg for eight weeks. In contrast, the saline-injected control group maintained an IOP of 11.10 ± 0.25 mm Hg (**Fig. 1F**). To quantify IOP elevation over time, we defined high IOP as persistently exceeding 15 mm Hg and measured its duration over the 8 weeks. The Viscobeads group exhibited high IOP for 97.2 ± 3.6% of the monitoring period, significantly longer than the saline group (1.1 ± 2.4%) (**Fig. 1G**). At 8 weeks post-injection, Viscobeads-injected eyes were comparable to or larger than saline-injected control eyes (**Fig. 1H**).

The TM consists of intertrabecular fenestrations with varying porosities, primarily composed of the uveal meshwork (UM), corneoscleral meshwork (CM), and juxtacanalicular tissue (JCT) ^[16]^. The UM features initial interlaminar spaces of approximately 25–27 μm, allowing the passage of larger beads. In contrast, smaller beads (0.2–2 μm) become trapped in the JCT layer, while mid-sized particles (2–15 μm) lodge within the CM layer ^[17]^. To examine the TM microarchitecture and surface morphology before and after Viscobeads injection, we performed scanning electron microscopy (SEM) (**Fig. 1I–J**). SEM images showed that the native TM has a porous morphology with filamentous structures (**Fig. 1I**, without injection). In contrast, after Viscobeads injection, the beads were observed adhering to the surface or embedded within the interlaminar spaces (**Fig. 1J**, with injection). Additionally, Micro- CT imaging of gel-embedded TM samples at 8 weeks post-injection confirmed the entrapment of Viscobeads within the tissue from multiple viewing angles, suggesting that Viscobeads remain in place long-term, potentially impairing aqueous outflow and contributing to glaucoma progression in mice (**Fig. 1K**).

### 2.2 Effect of Viscobeads Size Distribution on IOP Elevation

In the complex and dynamic fluidic environment of the AH, the transport of fluidic particles within the gradient porous structure of the trabecular meshwork is directly influenced by size distribution ^[9]^. To depict Viscobeads occlusion in the AH outflow pathway, we developed a simplified model featuring three layers-UM, CM and JCT- each characterized by pore sizes falling within a 1-20 μm range ^[17]^ (**Fig. 2A-E**). To mimic the natural size progression of TM structures, we established a discrete phase model (ANSYS Fluent simulation, **Fig. 2A-B**, and **Fig. S2**) and simulated the blocking effect of size-mixed spheres flowing through different layers. Heterogeneously distributed particles (Gaussian distribution with sizes ranging from 1 to 15 μm) were released from the inlet of a three-layered structure with gradient gap sizes to mimic the UM, CM, and JCT layers (10, 5, and 2 μm from inlet to outlet, respectively) in a discrete phase model. The blocking effect is visualized through the size distribution of trapped spheres and categorized into four planes: the inlet where the spheres are released, P0 (the initial point before larger gaps begin), P1 (the boundary between larger and medium gaps), P2 (the boundary between medium and small gaps), P3 (the plane beyond the smallest gaps), and the outlet (**Fig. 2B)**. The average sphere size changes from 7 μm at the inlet to 3.3 μm at the P3, indicating that the majority of the spheres were retained in the layered structures (**Fig. 2C**).

**Fig. 2:**
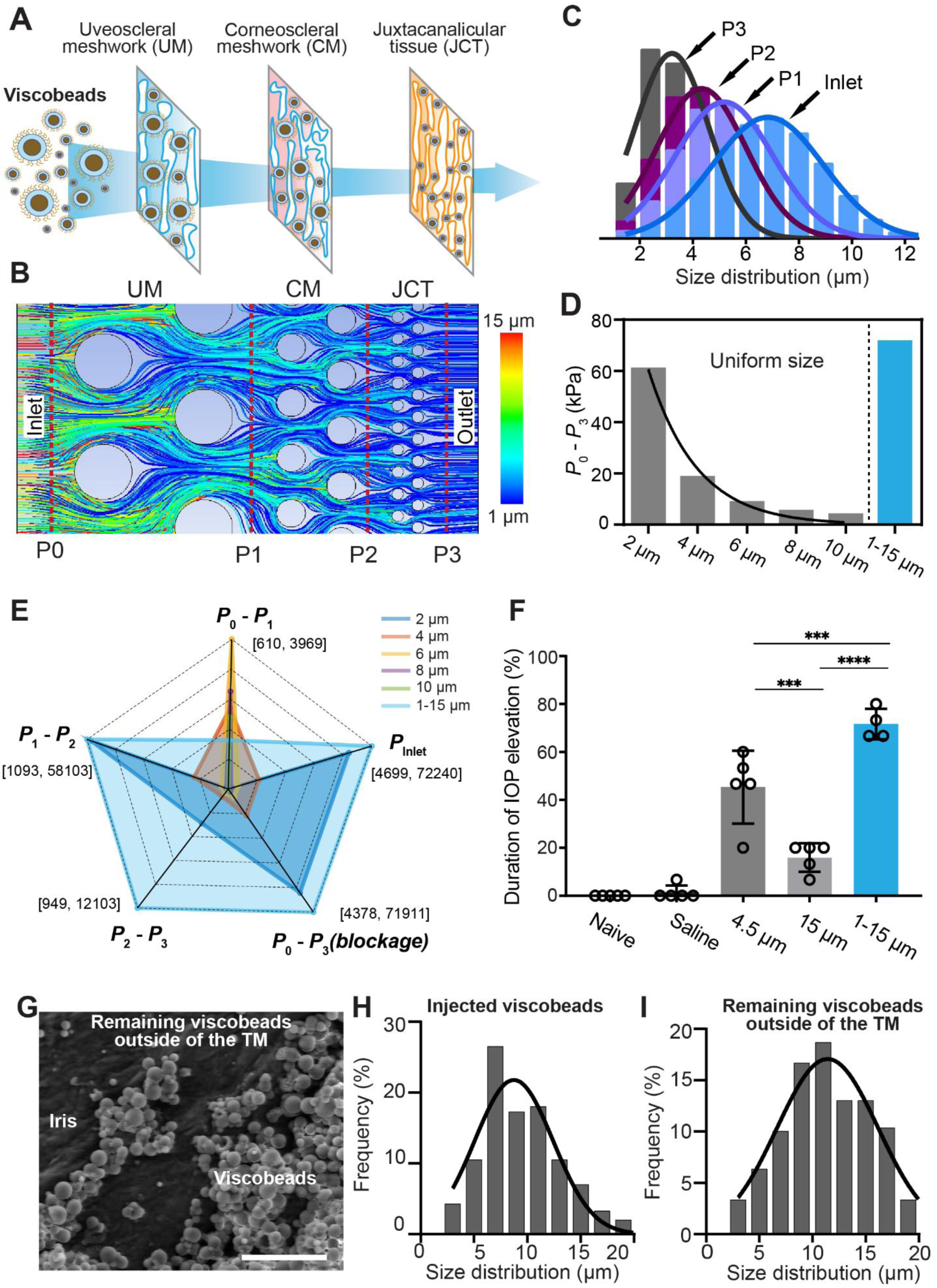
Effect of Viscobead Size Distribution on Modulating Aqueous Humor (AH) Outflow and IOP Elevation in Both Simulation and Experiment. (A) A simplified model illustrating Viscobeads blockage in the aqueous humor outflow pathway across three layers: UM, CM, and JCT. The initial interlaminar spaces in the UM measure approximately 25–27 μm, retaining larger Viscobeads. Smaller Viscobeads tend to clog in the JCT layer (0.2–2 μm), while mid-sized particles become trapped in the CM layer (2–15 μm). (B) Simulation of aqueous humor outflow obstruction using ANSYS Fluent. Heterogeneously distributed particles (Gaussian distribution, 1–15 μm) were released from the inlet of a three-layered structure with gradient gap sizes mimicking the UM, CM, and JCT layers (10, 5, and 2 μm from inlet to outlet, respectively) in a discrete phase model. Colored lines indicate trajectories of particles of different sizes. (C) Size distribution of particles at different layers (P0, P1, P2, and P3) in the ANSYS Fluent model, demonstrating particle entrapment at specific layers. (D) Simulation results showing pressure changes from P0 to P3 for both fixed-size beads (2, 4, 6, 8, and 10 μm) and heterogeneous beads (1–15 μm). (E) Radar plot of pressure changes across different layers from inlet to outlet, mimicking IOP, for fixed-size beads (2, 4, 6, 8, and 10 μm) and heterogeneous beads (1–15 μm). (F) Experimental data showing the duration of high IOP (≥15 mm Hg) in mice injected with 1–15 µm pure PS microbeads, 15 µm and 4.5 µm commercial microbeads, saline, and a naïve control group. Data are shown as mean ± SD. n=4 eyes for 1–15 µm pure PS; 5 eyes for other groups. One-way ANOVA with Tukey’s post hoc test (F(4,19) = 66.37, ****p<0.0001). (G) Representative SEM image showing remaining Viscobeads outside the trabecular meshwork at 8 weeks post-injection. The iris with residual Viscobeads was dissected and observed under SEM. Scale bar: 100 μm. (H-I) Size distribution of initially administered PS/PLGA Viscobeads (average: 9.61 ± 3.74 μm) and remaining Viscobeads outside the trabecular meshwork at 8 weeks post-injection (average: 11.90 ± 4.56 μm).

To assess the impact of bead size on IOP elevation, we compared pressure changes across different layers from inlet to outlet, mimicking IOP, for both fixed-size beads (2, 4, 6, 8, and 10 μm) and heterogeneous beads (1–15 μm). For size-fixed beads, the results showed a continuous decrease in pressure change as bead size increased, indicating that smaller beads more effectively obstruct aqueous humor outflow. Additionally, heterogeneous beads spanning the full-size range exhibited the highest pressure change from inlet to outlet, suggesting the most efficient blockage (**Fig. 2D**). Consistent with this, Goldmann-based analysis further predicts the highest relative IOP elevation under this condition (**Fig. S3**). To further evaluate the blocking effect of different bead sizes within specific TM layers, we analyzed pressure changes using a radar chart (**Fig. 2E**). The results indicated that larger beads (≥ 6 μm) caused greater pressure changes in the first layer with larger pores (P0-P1), whereas smaller beads (≤ 4 μm) were more effective at blocking smaller pores (P1-P2, P2-P3). Notably, heterogeneous beads demonstrated the most effective overall blockage (P0-P3). These simulation results suggest that heterogeneous beads combine the advantages of both larger and smaller beads, providing mechanical support from larger beads while enhancing obstruction through smaller beads.

To further evaluate the effect of microbead size on occlusion models, we synthesized pure polystyrene (PS) microbeads with a size distribution of 1–20 µm (**Supplementary method 2**) and compared the resulting IOP elevation in four groups of mice injected with fixed-size microbeads (4.5 μm and 15 μm) from commercially available PS beads. To avoid potential confounding effects from viscoelastic surfaces, we used PS, which is the same material used in commercial microbeads, without applying a PLGA coating. IOP monitoring over eight weeks revealed that uniform-sized commercial beads, whether 4.5 μm or 15 μm, resulted in lower IOP elevation (14.40 ± 0.26 mm Hg) and less stability compared to heterogeneous-size pure PS beads (19.08 ± 0.34 mm Hg) (**Fig. S4**). Over the 8-week monitoring period, the 1–20 μm pure PS microbead injection group exhibited the highest duration of IOP elevation among all treatment groups. The duration of high IOP in this group was significantly greater compared to both the Naive and saline control groups (p<0.0001). Specifically, the high IOP elevation in the 1-20 µm pure PS group reached 71.7 ± 3.2%, which was higher than in the commercial bead groups of 4.5 μm (45.3 ± 6.8%, p=0.0009) and the 15 μm group (16.0 ± 2.7%, p<0.0001). Additionally, while the 15 μm group showed a moderate increase in IOP compared to the Naive group (p=0.0373), its difference from the saline group was not statistically significant (p=0.0633). A significant difference was observed between the 4.5 μm and the 15 μm group (p=0.001) (**Fig. 2F**). These findings align with theoretical simulation data, indicating that smaller beads have a more effective blocking effect in fixed-size bead groups, while heterogeneous beads provide the highest IOP elevation.

In the context of a gradient porous structure, particles of different sizes become trapped according to local pore size. Larger pores accommodate larger beads, smaller beads infiltrate layers with smaller pores, and medium-sized particles become lodged within intermediate pore structures. This phenomenon is supported by both experimental data and theoretical modeling using the ANSYS Fluent simulation model (**Fig. 2A–F**). To determine which bead sizes were retained in the TM tissue, we dissected the mouse iris eight weeks post-injection and used SEM to observe the remaining beads on the iris surface (**Fig. 2G**), which were considered the non-trapped beads. We then compared their size distribution with the originally injected beads. The initial size distribution of injected beads was 9.6 ± 3.7 μm, whereas the non-trapped beads had an average size of 11.9 ± 4.6 μm (**Fig. 2H–I**). This observation suggests that smaller beads are more likely to be retained within TM tissue, contributing to enhanced blockage of aqueous humor outflow.

### 2.3 Biodegradable Surface Design for Chronic IOP Elevation

To improve the efficiency of AH flow blockage, we devised a strategy involving the incorporation of a biodegradable PLGA surface on the PS sphere cores. This design aimed to sustain elevated IOP by optimizing the retention of Viscobeads in TM tissues. In contrast to previously reported methods involving sequential injections of viscous substances followed by rigid polymer microbeads ^[8c, 12a]^, our approach leverages *in vivo* metabolic processes to degrade the surface PLGA materials, creating a viscoelastic contact between microbeads and tissues (**Fig. 3A, Fig. S5**). To mimic *in vivo* degradation conditions, we initially examined the degradation dynamics of PLGA *in vitro*. PLGA, a linear copolymer with adjustable constituent monomers, lactic (L) and glycolic (G) acid ^[18]^ (**Fig. 3B**), underwent incubation at 37 °C and 55 °C as an accelerated condition for 7 days. We measured the Raman spectra of PLGA glycolic and lactic units by comparing the intensities of featured peaks at 1452 and 1422 cm^−1^ ^[19]^ (**Fig. 3C**). In our design, the biodegradable PLGA serves as a viscous substance to promote adherence of the Viscobeads to the surrounding tissues, effectively enhancing accumulation of the beads in the TM tissue and improving aqueous humor obstruction (**Fig. 3D-E**). Through SEM observations, we confirmed that the microbeads became connected after incubation under *in vitro* conditions (**Fig. 3F**; 7-day incubation at 55 °C) and adhered to the surrounding tissues *in vivo* (**Fig. 3G**; 56 days post-injection). Additionally, neither the Viscobeads nor their degradation byproducts exhibited cytotoxicity compared to the PBS control group, as assessed using in vitro cultured cells following an established mouse corneal tissue culture protocol ^[20]^ (**Supplementary method 4, Fig. S6 A-B**). We further evaluated cellular biocompatibility by comparing Viscobeads with commercially available PS beads and observed no significant differences in cell viability for both trabecular meshwork cells and Neuro-2A cells (**Fig. S6 C-D**).

**Fig. 3:**
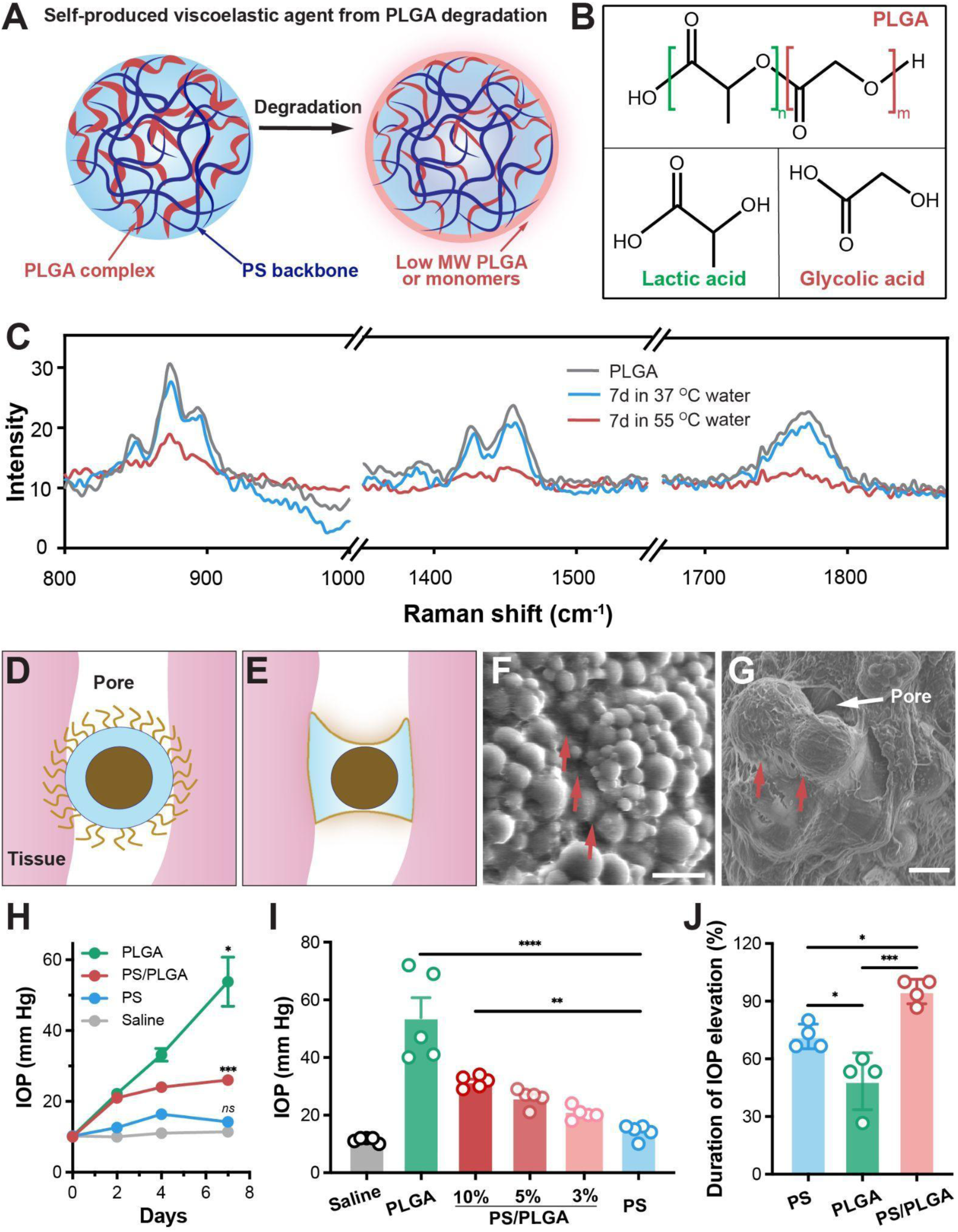
The Mechanism of Self-Production of Viscoelastic Agents Through Degradation. (A) PLGA degrades into low molecular weight PLGA or monomers, which serve as viscoelastic materials. (B) PLGA is a linear copolymer composed of tunable constituent monomers, lactic acid (L) and glycolic acid (G). (C) Raman spectra of PLGA Viscobeads before and after submersion for 7 days at 37 °C and 55 °C. (D-E) The illustration demonstrates that the biodegradable PLGA acts as a viscous substance, efficiently binding the Viscobeads to the surrounding tissue, thereby augmenting the entrapment of beads in TM tissue and contributing to effective aqueous humor blockage. (F-G) Representative SEM images show Viscobeads immersed in a 55°C water bath for 7 days in the in vitro condition (F) and Viscobeads entrapped on the surface pores of the trabecular meshwork (G). The red arrow indicates the degraded PLGA viscous substance. Scale bar: 20 µm for (F) and 10 µm for (G). (H) The IOP during the first week in mice injected with different microbead compositions. n=5 eyes. Two-way repeated measures ANOVA with Geisser-Greenhouse’s correction (ε=0.4199) and Tukey’s post hoc test (Time x Groups F(9, 48)=22.62, ****p<0.0001, Time F(1.260, 20.16)=61.05, ****p<0.0001, Groups F(3, 16)=51.94, ****p<0.0001, Subject F(16, 48)=1.814, p=0.0570). (I) Comparison of IOP elevation levels during the first week post-injection of saline, PLGA, PS, and PS/PLGA Viscobeads containing 10%, 5%, and 3% PLGA components (w/w). n=5 eyes. One-way ANOVA with Tukey’s post hoc test (F(5, 24) = 26.76, ****p<0.0001). (J) Duration of high IOP (≥15 mm Hg) over an 8-week evaluation period in mice injected with 1–20 µm pure PS microbeads, pure PLGA microbeads, and PS/PLGA Viscobeads. n=4 eyes. One-way ANOVA with Tukey’s post hoc test (F(2, 9) = 21.78, ***p= 0.0004).

Manipulating the composition can create distinct mechanical properties and subsequently impact IOP elevation dynamics in vivo. We prepared microbeads with a similar size distribution but different polymer compositions: pure PS, pure PLGA, and PS/PLGA Viscobeads. Three groups of mice were injected with PS, PLGA and PS/PLGA Viscobeads into the anterior chambers of mice, respectively, while the control group of mice was injected with the same volume of saline. Daily IOP measurements were recorded in all groups over a 7-day post-injection period. Following the rapid degradation of PLGA, both the groups of mice injected with pure PLGA microbeads and PS/PLGA Viscobeads exhibited a rapid elevation in IOP two days post-injection (**Fig. 3H**). The PS group showed no significant difference in IOP compared to the saline group (p=0.2015) on day 7. In contrast, the PS/PLGA group exhibited a significant increase in IOP starting day 2 (p=0.004), reaching a higher IOP level (≥ 15 mm Hg) and maintaining it throughout the week (on day 7, p=0.0007). Meanwhile, the PLGA group experienced the highest IOP elevation, peaking during the week (p=0.0123). Due to the ongoing degradation of PLGA and the generation of viscous substances, the pure PLGA beads showed a dramatic increase in IOP to 53.8 ± 6.93 mm Hg within one week, but the duration of maintaining high IOP was notably lower than PS or PS/PLGA Viscobeads group (PLGA *vs* PS p<0.0001; 10% PS/PLGA *vs* PS p=0.0003) (**Fig. 3I**). In other rodent models, such as rats, it has also been reported that PLGA microspheres have been used to elevate IOP to 15–30 mm Hg for 8–24 weeks with a single injection ^[21]^. The IOP elevation in the pure PS group was much lower than that in the PLGA and PS/PLGA Viscobeads groups. Unlike the acute and severe blocking effect of pure PLGA microbeads or the insufficient blocking effect of pure PS microbeads, the PS/PLGA Viscobeads group exhibited sufficient IOP elevation while maintaining it in a relatively stable high IOP range over time (26.0 ± 1.34 mm Hg on day 7).

The elevated and stable IOP characteristic of the PS/PLGA hybrid Viscobeads can be finely adjusted by manipulating the PLGA mass ratios because of the *in vivo* degradation dynamics of PLGA. We synthesized a series of PS/PLGA Viscobeads with PLGA mass ratios of 3%, 5%, and 10%, resulting in varying IOP levels within the range of 20.6 mm Hg to 31.6 mm Hg (**Fig. 3I**). Under the conditions of pure PS or pure PLGA, the IOP exhibited either insufficient elevation amplitude or duration for maintaining high IOP. The pure PS beads group reached 14.2 mm Hg, which was only 2.8 mm Hg higher than the saline group (11.4 mm Hg). Conversely, the pure PLGA beads achieved a severe acute elevation to 53.8 mm Hg but could sustain elevation for 48.3% of the observation period (PS *vs* PLGA p=0.0227) (**Fig. 3I-J**). Only the PS/PLGA Viscobeads demonstrated both effective IOP elevation (>15 mm Hg) and maintained 95% of the duration within the high IOP range during an additional 8-week long-term evaluation (PS *vs* PS/PLGA p=0.0227; PLGA *vs* PS/PLGA p=0.0003). We also observed significant variation in IOP elevation for the pure PLGA and pure PS bead groups across different study batches.

### 2.4 Chronic High IOP-Induced Neurodegeneration

We then assessed the chronic ocular hypertensive response and glaucomatous neurodegeneration by examining post-mortem retina tissues from mice injected with Viscobeads. To evaluate the progression of glaucomatous neurodegeneration, we quantified the RGC survival rate at different time points post-Viscobeads injection. The somatic structure of surviving RGC was labeled with RNA-binding protein with multiple splicing (RBPMS) antibodies on retinal wholemounts ^[22]^ (**Fig. 4A**) . To elucidate the impact of sustained high IOP on RGC death, we calculated the percentages of RBPMS-labeled RGC from retinal tissues at a series of specified time points and compared them to retinal tissues dissected on the same day post-injection (**Fig. 4B**). Notably, we observed a significant decrease in RGC survival percentages over an 8-week observation period. Among the eye tissues injected with Viscobeads, the RGC survival rate decreased to 84.48 ± 2.22% at week 4 (p=0.0017) and further declined to 66.43 ± 2.63% at 8 weeks post-injection (p<0.0001).

**Fig. 4:**
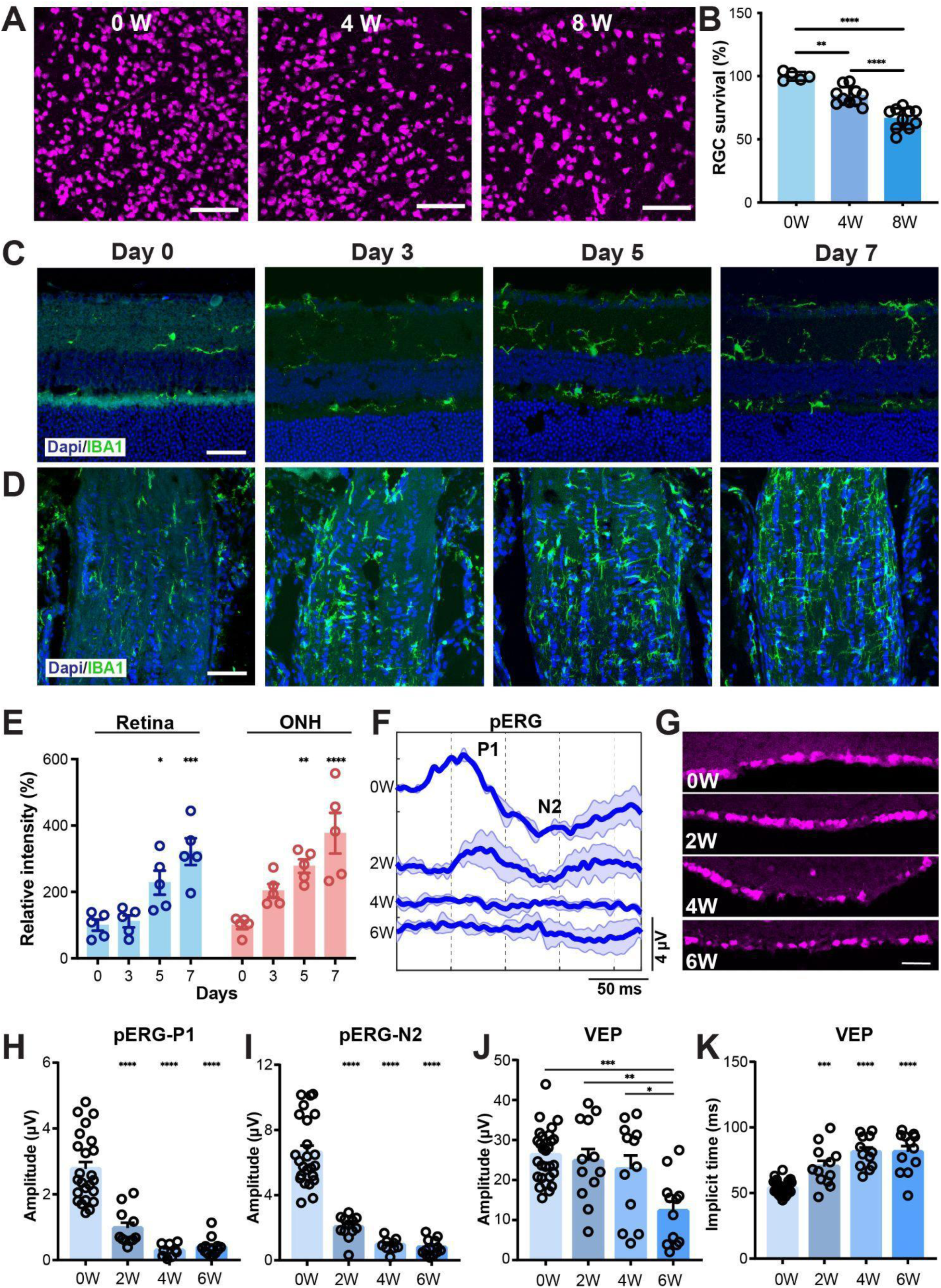
Glaucomatous Neurodegeneration in the Viscobeads-Induced Ocular Hypertension Model. (A) Representative whole-mount retinal confocal images from mice at 0, 4, and 8 weeks after PS/PLGA Viscobeads injection. RBPMS antibody was used for immunohistochemical staining of RGCs. Scale bars: 100 μm. (B) Quantification of RGC survival at different time points after PS/PLGA Viscobeads injection. Data are shown as mean ± SD. n=5 eyes for 0W; 10 eyes for 4W and 8W. One-way measures ANOVA with Tukey’s post hoc test (F(2, 22)=39.98, ****p<0.0001). (C-E) Representative confocal images (C-D) and quantification (E) of retinal cryosections (C) and optic nerve head (D) stained with DAPI and IBA1 at days 0, 3, 5, and 7 post-injections. Scale bars: 50 μm. Data are shown as mean ± S.E.M. n=5 eyes. Two-way repeated measures ANOVA with Tukey’s post hoc test (Time x Intensity F(3, 24)=0.8545, p=0.4780, Time F(3, 24)=28.67, ****p<0.0001, Intensity F(1, 8)=3.026, p=0.1202, Subject F(8, 24)=1.917, p=0.1040). (F-K) Electrophysiological assessments using pattern electroretinogram (pERG) and flash visual evoked potential (VEP) to evaluate RGC function and visual pathway integrity during glaucoma progression. (F) Representative pERG signals in mice from week 0 (pre-injection) to week 6 post-injection. (G) Representative confocal images of retinal sections. Scale bar: 50 μm. (H-I) Statistical quantification of the amplitudes of P1 and N1 of the pERG signals before and after Viscobeads injection. n=24 eyes for 0W; 12 eyes for 2W and 6W; 9 recordings for 4W. One-way measures ANOVA with Tukey’s post hoc test (pERG-P1: F(3, 52)=42.44, ****p<0.0001; pERG-N2: F(3, 53)= 65.68, ****p<0.0001). (J-K) Statistical quantification of the amplitude (J) and implicit time (K) of the VEP signals before and after Viscobeads injection. n=28 eyes for 0W; 12 eyes for other groups. One-way repeated measures ANOVA with Tukey’s post hoc test (Amplitude: F(3, 60)=7.292, ***p=0.0003; Implicit time: F(3, 60)=25.83, ****p<0.0001). Data are shown as mean ± S.E.M.

As an early event in the progression of glaucoma, microglia undergo morphological changes, proliferation, and migration into the injury site ^[23]^. Microglial cells produce proinflammatory cytokines and increase oxidative and nitrification reactions, exerting further negative effects on retinal neurons ^[23]^. To investigate the spatial and temporal dynamics of the activated microglia during glaucomatous progression, we examined microglia and macrophages by immunolabeling the retina and optic nerve head (ONH) using ionized calcium-binding adaptor molecule 1 (IBA1) antibody (**Fig. 4C-D**). Concurrent with the chronic high IOP induced by the Viscobeads, the intensity of IBA1+ cells in retinal and ONH tissues over time. In the retina, fluorescence intensity of the IBA1+ cells showed an upward trend from day 3, with a notable rise on day 5 (p=0.0411) and a significant increase on day 7 (p=0.0002). In the ONH, intensity increased from baseline on day 5 (p=0.0012) and continued to rise, exceeding three times the baseline level on day 7 (p<0.0001) (**Fig. 4E**). Similar trends were observed in activated microglia labeled with CD68 in both retinal and ONH tissues at the early stages of ocular hypertension (**Fig. S7**).

### 2.5 Electrophysiological Assessments of Changes in RGC Function and Visual Pathway Integrity During Glaucoma Progression

To monitor retinal function post-Viscobeads injection, we next employed non-invasive pattern electroretinogram (pERG) recordings. pERG is a crucial electrophysiological assessment of general RGC function (**Fig. 4F-I**). In this test, the ERG responses were stimulated by contrast-reversing horizontal bars that alternate at a constant mean luminance. pERG measurements included the mean amplitude of the P1 and N2 components. The P1 component represents the initial positive deflection, originating from both RGC and outer retinal photoreceptor cells. The mean P1 amplitude was measured as the difference between the average nadir of the N1 component and the average peak of the P1 component. The N2 component is the negative deflection that follows the P1 component, originating from the inner retina and reflecting RGC function. The mean N2 amplitude was calculated as the difference between the average peak of the P1 component and the average nadir of the N2 component (**Fig. 4F**). At week 2 post-Viscobeads injection, both the P1 and N2 components were significantly decreased compared to the control group with PBS injection, from 2.77 ± 0.21 μV to 0.97 ± 0.16 μV for P1 and from 6.62 ± 0.42 μV to 2.04 ± 0.20 μV for N2. This decrease continued to 0.40 ± 0.08 μV for P1 and 0.80 ± 0.14 μV for N2 at week 6 post-Viscobeads injection (0 week *vs* 2 week, 4 week and 6 week p<0.0001) (**Fig. 4H-I**). These results suggest that RGC function diminishes along with glaucoma progression, even at an early stage where RGC does not show a significant decrease based on RBPMS staining (**Fig. 4G**).

We then used flash visual evoked potential (VEP) to evaluate retinal and cortical function, as well as the integrity of the visual pathway from the retina to the cortex. This approach helps identify optic nerve fiber loss and glaucomatous damage. For the flash VEP recording, a flickering light stimulus (duration: 5 ms, frequency: 0.1 Hz, light intensity: 1 cd s⁻² m⁻²) was applied to one eye while recording the cortical responses. The average response to the stimulus was obtained for subsequent analysis. In VEP measurements, we calculated the amplitude and implicit time for quantitative analysis. The VEP amplitude (**Fig. 4J**) was determined as the difference between the baseline and first negative peak within 300 ms after stimulus presentation, reflecting the strength of the signal reaching the visual cortex and correlating with the number of functional RGC. The implicit time (**Fig. 4K**) was defined as the interval from the onset of the visual stimulus to the point where the first negative peak is reached, which represents the conduction time it takes for the signal to be transmitted from the retina to the visual cortex. The average amplitude in the control group was 26.25 ± 1.25 μV. In mice at 2, 4, and 6 weeks post-Viscobeads injection, the average amplitudes were 24.83 ± 2.95 μV, 22.69 ± 3.47 μV, and 12.26 ± 2.45 μV, respectively (0W *vs* 6W p=0.0001; 2W *vs* 6W p=0.0051; 4W *vs* 6W p=0.0271) (**Fig. 4J**). Endpoint waveform comparisons revealed a marked reduction in amplitude in Viscobeads-injected mice. The average implicit time in the control group was 53.84 ± 1.05 ms. In contrast, mice at 2, 4, and 6 weeks post-Viscobeads injection exhibited significantly prolonged implicit times of 70.17 ± 4.39 ms, 81.21 ± 3.47 ms, and 81.29 ± 4.86 ms, respectively (0W *vs* 2W p=0.0005; 0W *vs* 4W and 6W p<0.0001) (**Fig. 4K**). As the duration of Viscobeads injection increased, the amplitude of the VEP gradually decreased, suggesting a weakening of signal transmission, indicating that RGC gradually experienced damage and death. Additionally, the prolonged implicit time may indicate the progression of glaucomatous optic nerve damage. These findings suggest that the integrity of the visual pathway is progressively compromised during glaucoma progression.

### 2.6 A Viscobeads-Enabled Experimental Glaucoma Model for Gene Therapy Development

The reliability of sustaining chronic high IOP elevation and neurodegeneration supports a dependable and accessible model for the testing of therapeutic candidates. As a proof-of-concept for gene therapy development, we applied the Viscobeads experimental glaucoma mouse model and tested genetic approaches to mitigate glaucomatous neurodegeneration. Gene candidates were initially selected based on their demonstrated neuroprotective effects in other axotomy animal models, such as optic nerve crush, as established in our prior research and by others ^[13, 24]^. We chose to test the following genes: phosphatase and tensin homolog (PTEN), a tumor-suppressor; activating transcription factor 3 (ATF3), a transcription factor; C/EBP homologous protein (CHOP), a transcription factor responsive to endoplasmic reticulum (ER) stress; doublecortin-like kinase 2 (DCLK2), a key regulator of growth cone formation and axon regeneration; armadillo repeat-containing X-linked protein 1 (ARMCX1), a regulator of mitochondrial transport; Lin-28 homologue A (LIN28), an RNA-binding protein; and a combination of LIN28 and insulin-like growth factor 1 (IGF-1). We classified the gene candidates into two groups, knockout and overexpression, based on their functional roles. For the knockout group, we designed single-guide RNAs (sgRNAs) targeting PTEN, ATF3, and CHOP, following established protocols. These sgRNAs were packaged into adeno-associated viruses (AAV) and delivered into the vitreous bodies of C57BL/6J wild-type mouse eyes via co-injection of AAV2-sgRNA and AAV2-Cas9. For the overexpression group, we packaged the selected genes, including DCLK2, ARMCX1, LIN28, and IGF-1, into AAV and introduced them into the vitreous bodies of C57BL/6J wild-type mouse eyes. Two weeks following these viral infections, PS/PLGA Viscobeads were intracamerally injected into the anterior chamber to induce ocular hypertension (**Fig. 5A**).

**Fig. 5:**
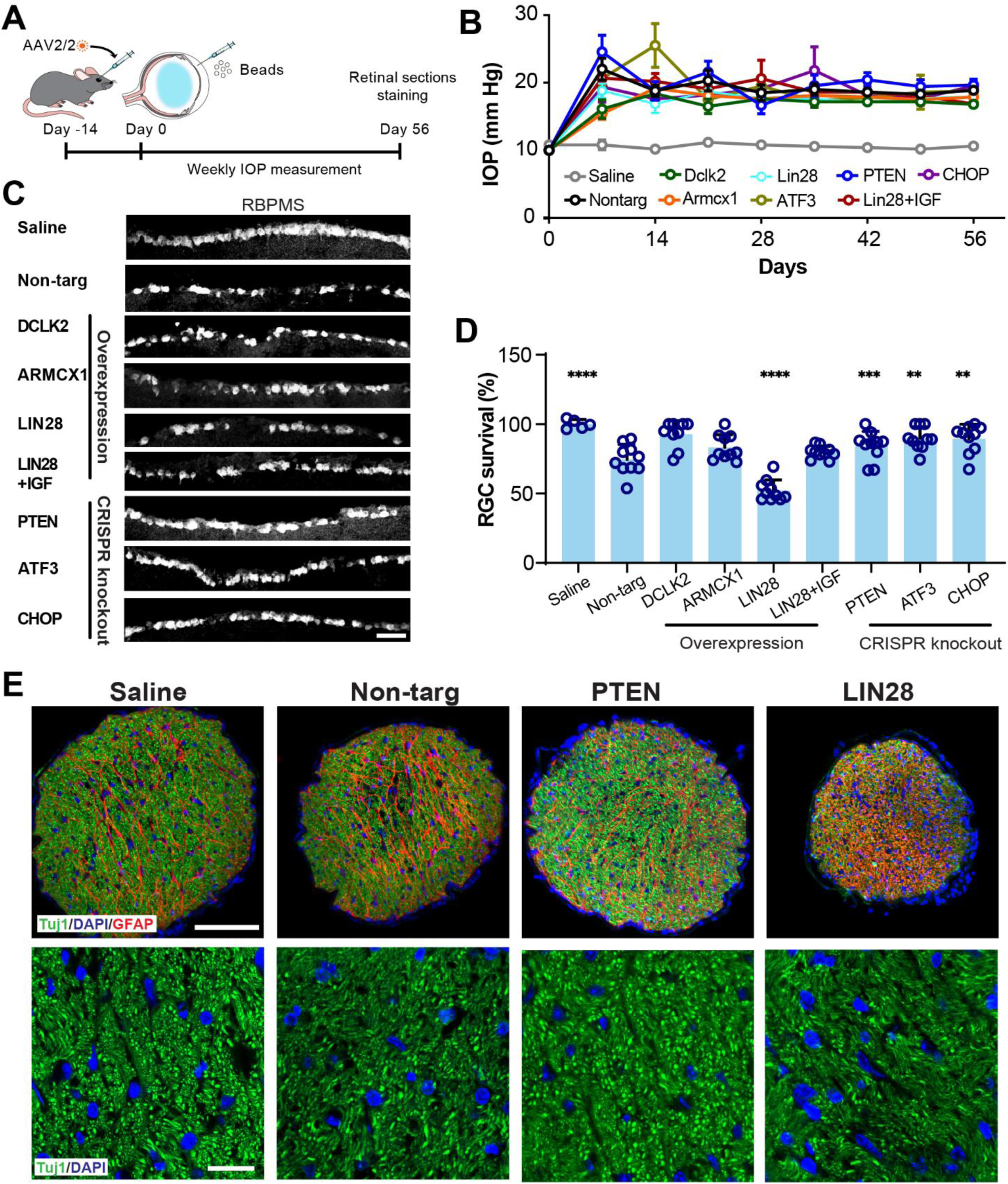
Genetic Manipulation in the Retina to Prevent RGC Loss in the Viscobead-Induced Glaucoma Model. (A) Schematic illustration of AAV2 vector delivery via intravitreal injection, administered two weeks before microbead injection on day 0. For the DCLK2, ARMCX1, LIN28, and IGF-1 overexpression groups, AAV2 carrying the respective cDNAs was injected into the vitreous body of C57BL/6J wild-type mouse eyes on day -14. For the PTEN, ATF3, and CHOP knockout groups, AAV2-sgRNAs and AAV2-Cas9 were delivered intravitreally into C57BL/6J wild-type mouse eyes on day -14. (B) 8-week time-course of IOP changes across different groups of mice. (C) Representative images of retinal sections showing RGC survival with AAV injections. Scale bar: 20 μm. (D) Quantification of RGC survival after 8 weeks of PS/PLGA Viscobeads injection. One-way measures ANOVA with Dunnett’s multiple comparison test with the non-target sgRNA group as the reference (non-target *vs* other groups; F(8, 75) = 21.26 ****p<0.0001). (E) Representative confocal images of optic nerve cross-sections from mice with saline injection or non-targeting sgRNA, PTEN, and LIN28 AAV injection. Scale bar for the upper panel: 100 μm; bottom panels: 20 μm. Data are shown as mean ± SD. n=5 eyes for Saline; 10 eyes for other groups.

Over an 8-week observation, all mice consistently exhibited elevated IOP (**Fig. 5B**), validating the reliability of Viscobeads as an experimental glaucoma model and its compatibility with viral vector introduction. Among the selected gene candidates, we observed enhanced RGC survival rates, which indicate improved RGC protection, in the mice via CRISPR-mediated knockout of PTEN (p=0.0002), ATF3 (p=0.0019), and CHOP (p=0.0024) relative to the non-targeting sgRNA. However, mice with LIN28 overexpression demonstrated a dramatic reduction in survival rates after 8 weeks of Viscobeads injection (p<0.0001). Mice overexpressing ARMCX1 (p=0.1761), DCLK2 (p=0.1031), or the LIN28+IGF combination (p=0.7136) showed no significant differences in RGC survival rates when compared to the control mice that received non-targeting sgRNA injections (**Fig. 5C-D**). Upon examining retinal sections with immunohistology staining of RGC axons and glial cells, we identified similar protective effects of PTEN, ATF3, and CHOP, as well as an adverse effect of LIN28 on RGC axon density through immunohistochemistry of optic nerve cross-sections (**Fig. 5E**).

### 2.7 Sleep Behavior Changes in Experimental Glaucoma Mouse Models

Inspired by the clinical observation of sleep behavior changes in human patients with glaucoma, we applied an experimental glaucoma mouse model to investigate the relationship between sleep disturbances and glaucoma. To preserve natural behaviors such as breathing rhythms and sleep patterns from untethered mice, we employed a non-invasive PiezoSleep monitoring system, which utilizes a piezoelectric recording pad placed beneath the mouse cage to capture and analyze sleep behaviors ^[25]^ (**Fig. 6A-B**). Unlike traditional electrophysiological recording methods that require surgical procedures or tethered setups, this approach enables the undisturbed assessment of sleep behaviors in freely behaving mice with more than 90% accuracy ^[25–26]^. All mice were continuously monitored for over 36 hours, and the recordings were trimmed to a standardized 24-hour period, encompassing a full 12-hour light and 12-hour dark cycle (**Fig. 6C**).

**Fig. 6:**
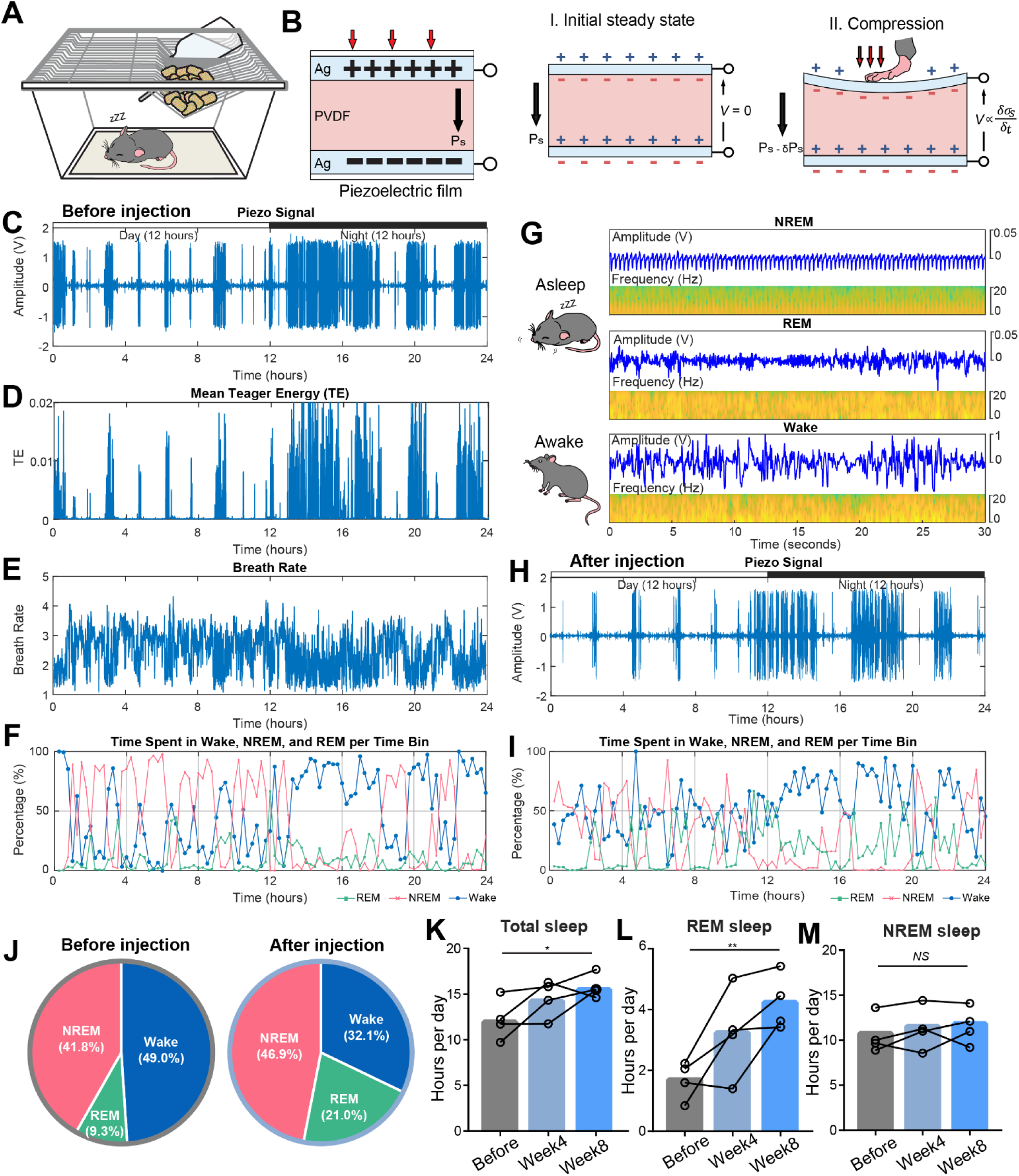
Non-Invasive Evaluation of Sleep Pattern Changes in Experimental Glaucoma Mouse Models. (A-B) Schematic of the non-invasive PiezoSleep mouse monitoring system, which utilizes a piezoelectric pad to detect sleep/wake status based on movement-related features, as movement induces voltage changes in the piezoelectric pad. (C) Representative piezoelectric signal output over a 24-hour period for a control mouse under a 12-hour light/12-hour dark cycle. (D-E) Sleep stage classification based on published features: (D) Teager energy (TE) quantifies motion, (E) breathing rate, and breathing regularity or variability (BRV). (F) Wakefulness is characterized by high TE, rapid eye movement (REM) sleep by inactivity with irregular breathing, and non-rapid eye movement (NREM) sleep by inactivity with regular breathing. (G) Summary of classification criteria: wakefulness corresponds to high activity (TE), REM sleep to inactivity with irregular breathing, and NREM sleep to inactivity with regular breathing. (H-I) Representative piezoelectric signal output for the same mouse following Viscobead injection (H) and its corresponding classification into wakefulness, REM sleep, and NREM sleep over 24 hours (I). (J) The pie chart shows the percentage of time spent in wakefulness, REM sleep, and NREM sleep before and after injection in the same mouse. (K-M) Total sleep, REM sleep, and NREM sleep duration per 24 hours in mice before injection, and at 4 and 8 weeks post-injection. n=4 mice. One-way repeated measures ANOVA with Dunnett’s post hoc test (Total sleep: Time F(1.697, 5.092) = 6.176, *p=0.0460; Mice F(3, 6) = 3.143, p = 0.1081, REM: Time F(1.057, 3.172) = 12.91, *p=0.0334; Mice F(3,6) = 4.694, p=0.0514, NREM sleep: Time F(1.738, 5.215) = 0.8447, p=0.4653; Mice F(3, 6) = 8.308, **p=0.0148).

Using previously established features that quantify motion (Teager energy, TE), breathing rate (f), and breathing regularity or variability (BRV) ^[25–26]^ (**Fig. 6D-E**), we classified wakefulness, non-rapid eye movement (NREM), and rapid eye movement (REM) sleep stages (**Fig. 6F**). A low-frequency (0.5 Hz to 18 Hz) band-pass filter was applied to minimize noise and artifacts. Wakefulness was defined by high activity (TE), REM sleep by inactivity with irregular breathing, and NREM sleep by inactivity with regular breathing (**Fig. 6G**). Our results showed that the overall sleep and wake patterns in mice before and after injection remained largely unchanged: all mice exhibited greater activity during nighttime and increased sleep duration during daytime (**Fig. 6H-I**). However, the proportion of REM and NREM sleep showed differences (**Fig. 6J**). Before injection, the total sleep duration per 24 hours was 12.2 ± 2.3 hours in the Viscobeads group and 12.9 ± 1.5 hours in the saline group. Following the Viscobeads injection, total sleep duration increased to 14.6 ± 2.0 hours at four weeks post-injection and 15.8 ± 1.3 hours at eight weeks post-injection. In contrast, saline-injected mice showed no significant change in total sleep duration, recording 13.7 ± 0.6 hours at four weeks and 12.8 ± 2.6 hours at eight weeks post-injection (Before *vs* Week4 p=0.222, Before *vs* Week8 p=0.0328). Notably, Viscobeads-injected mice exhibited increased REM sleep, particularly at eight weeks post-injection. The total REM sleep duration increased from 1.7 ± 0.6 hours before injection to 3.2 ± 1.5 hours at four weeks and 4.2 ± 0.9 hours at eight weeks post-injection (Before *vs* Week4 p=0.1749, Before *vs* Week8 p=0.0032), while NREM sleep duration remained unchanged (Before *vs* Week4 p=0.5007, Before *vs* Week8 p=0.4537) (**Fig. 6K-M, Fig. S8**). In contrast, mice injected with saline exhibited stable sleep durations across different recording sessions and among different mice housed on the same recording platform (**Fig. S8**).

## 3 Discussion

The Viscobeads technology provides a bio-interface solution to manipulate fluidic transport in heterogeneously structured biological tissues and can be directly applied as an injectable method to create an experimental glaucoma mouse model with sustained intraocular hypertension. The “hard” polymer core provides mechanical support for long-term retention while the biodegradable “soft” surface provides viscoelastic contact to tissues, yielding composite viscoelastic particles. Tuning the composition of the hybrid materials can affect the surface materials’ biodegradation dynamics and further regulate the occlusion effect. Instead of using uniform-sized microbeads, the Viscobeads (1-20 μm) were designed to match the size gradient of porous structures in the trabecular meshwork that regulate intraocular pressure. The combination of hybrid material composition and a precise size distribution optimized the beads-tissue infiltration after a single-dose and produced long-term bead retention in the TM. IOP elevation (21.4 ± 1.39 mm Hg) was subsequently sustained over 8 weeks, corresponding to observed glaucomatous neurodegeneration. Over 34% RGC death was observed along with a dramatic oscillatory potential change in electroretinography. While analyzing pERG and VEP signals and retinal tissues, we found that RGC function is impaired at an early stage of ocular hypertension, even before observable RGC loss in tissues. In the later stages of glaucoma, the integrity of the visual pathway is likely affected in this experimental model.

The Viscobeads design is grounded in the *in vivo* aqueous humor flow dynamics and graded architecture of the trabecular meshwork. Rather than producing primarily surface-level occlusion with uniform-sized beads, heterogeneously sized beads matched to the trabecular meshwork pore distribution create multilayer obstruction throughout the tissue. The viscoelastic surfaces of the hybrid beads further promote prolonged retention under dynamic fluid-flow conditions. Both the size distribution and degradation kinetics of Viscobeads can be tuned during synthesis, providing a versatile engineering strategy for adapting the occlusion model to different tissue architectures. As the PLGA progressively degrades, early softening may enhance bead–bead and bead–tissue adhesion and transiently strengthen occlusion, whereas continued erosion at later stages may reduce interfacial retention and partially restore aqueous humor outflow, while the non-degradable PS cores maintain residual mechanical obstruction. The non-degradable PS cores are expected to remain within the iridocorneal angle and trabecular meshwork and may persist long term. Accordingly, the Viscobeads platform is intended solely for experimental disease modeling rather than therapeutic or clinical implantation.

To model glaucoma-associated IOP elevation, several engineered occlusion strategies have been developed to impede aqueous humor outflow through the conventional TM pathway, including conventional microbead occlusion (polystyrene/latex),^[11]^ magnetic microbead-assisted occlusion,^[6c, 27]^ viscoelastic (e.g., hyaluronic acid) injection,^[28]^ and silicone oil–induced obstruction.^[29]^ These approaches typically elevate IOP to approximately 20– 30 mm Hg and result in 10–70% RGC loss in mice, depending on the model and duration. However, they often exhibit substantial variability in IOP magnitude and duration due to inconsistent material retention, heterogeneous outflow blockage, and differences in surgical procedures. In contrast, the Viscobeads system enables more stable and tunable IOP elevation by improving bead-tissue interactions within the trabecular meshwork, thereby reducing experimental variability and providing a more reliable platform for mechanistic and therapeutic studies.

The Viscobeads experimental glaucoma model provides a reliable *in vivo* testbed that replicates glaucoma-like high IOP and retinal degeneration. Introducing the Viscobeads is compatible with other surgical procedures, such as injecting viral vectors to manipulate gene expression. In a proof-of-concept application, we used the Viscobeads glaucoma model to assay a series of previously selected gene candidates and gained new insights into the neuroprotective effects of these genes. Notably, during this screening, we identified that LIN28 exhibited detrimental effects on RGC survival. Our findings challenge the widely held belief that the optic nerve crush model sufficiently mimics glaucoma neurodegeneration. The distinct results for LIN28 between the optic nerve crush model^[30]^ and our experimental glaucoma model underscore our model’s reliability and its potential to provide more accurate insights into glaucoma pathology.

Additionally, the Viscobeads-induced experimental glaucoma model provides a platform for tracking longitudinal changes in sleep behavior during disease progression. When combined with a non-invasive sleep monitoring system, this approach enables continuous assessment of sleep-related disturbances with minimal experimental interference, thereby offering insight into the broader systemic consequences of glaucoma. Although this mouse model does not reproduce all clinical features of human glaucoma quantitatively, it captures several key disease-associated changes in parallel. The IOP elevation observed in mice reflects the same pathogenic driver central to many forms of human glaucoma, even though the absolute IOP levels, temporal profile, and experimental induction differ from those seen in patients. Likewise, the RGC loss detected in this model recapitulates the core neurodegenerative feature of glaucoma, although the extent, regional distribution, and rate of progression are more controlled and compressed than in the heterogeneous clinical setting. With respect to behavior, the altered sleep pattern observed in mice is not intended to mirror the full complexity of sleep dysfunction in glaucoma patients, but rather to model a related disruption of retinal-brain signaling. In humans, glaucoma has been associated with sleep and circadian abnormalities, and these effects have been linked in part to dysfunction or loss of intrinsically photosensitive retinal ganglion cells (ipRGCs), which contribute to non-image-forming visual functions such as circadian entrainment and sleep regulation^[31]^. In this context, the sleep phenotype detected in the Viscobeads model supports its translational relevance while also highlighting that the underlying mechanisms remain incompletely understood. This model provides a valuable *in vivo* framework for integrating ocular hypertension, glaucomatous retinal injury, and behavioral outcomes to investigate how glaucoma progression may influence sleep regulation.

Overall, the Viscobeads technique offers a new perspective on the materials-tissue interfaces used to regulate fluid transport *in vivo*. Similar interface designs could be further extended to other biological tissues that regulate biofluid flow with micro-architectures, such as the lymphatic system, pulmonary alveolus, and bones ^[32]^. As a tool to study the neurodegeneration of glaucoma, the Viscobeads technique provides a scalable and accessible engineering method to introduce *in vivo* occlusion for models of ocular hypertension. The injection is minimally invasive and compatible with the delivery of other genetic or pharmaceutical reagents. By sustaining a prolonged and stable high IOP *in vivo* condition upon a single injection, the Viscobeads technique provides a reliable testbed for future therapeutic intervention development.

## 4 Conclusion

In conclusion, we developed a biologically driven Viscobeads platform that improves bead-tissue interfacing within the trabecular meshwork and enables stable, long-term ocular hypertension after a single injection. By combining a structural PS core with a viscoelastic PLGA shell, this composite design better accommodates the graded porous architecture of the outflow pathway and produces sustained IOP elevation sufficient to drive progressive glaucomatous pathology. The model recapitulates key features of glaucoma progression, including early retinal ganglion cell dysfunction, later visual pathway impairment, and substantial RGC loss, while also providing a practical *in vivo* platform for genetic screening and broader phenotypic assessment. Using this system, we identified neuroprotective and detrimental gene targets for RGC survival and further revealed sleep-related behavioral changes associated with ocular hypertension. Together, these results establish Viscobeads as a versatile and accessible materials-based strategy for modeling chronic glaucoma and studying its neural and systemic consequences, with potential utility for evaluating future genetic and pharmacological therapies.

## 5 Materials and Methods

### 5.1 PS/PLGA Viscobeads fabrication

Detailed fabrication procedures are provided in Supplementary Methods. Briefly, PS/PLGA Viscobeads were prepared using a single emulsion method in an oil-in-water configuration. To prepare the PS microspheres, 1 mL of deionized (DI) water was combined with 3 mL of a 0.1 g/mL PS in chloroform solution. For visualization in confocal microscopy, rhodamine B (83689, Sigma) aqueous solution was used instead of DI water. This mixture was then emulsified using a high-speed homogenizer at 5000 rpm for 30 seconds. The resulting primary emulsion was promptly poured into 15 mL of a 2% PVA aqueous solution and stirred to create a double emulsion. To remove the organic solvents, the mixture was transferred into 100 mL of 55°C water and stirred for 30 minutes. The resulting PS microspheres were subsequently washed with DI water three times and centrifuged at 2000 rpm. For surface coating, the dried PS microspheres were introduced into 3 mL of a 0.02 g/mL PLGA solution in dichloromethane. After adding 15 mL of a 2% PVA aqueous solution, the mixture was emulsified at 5000 rpm for 30 seconds. Following the evaporation of organic solvents, PS/PLGA Viscobeads were centrifuged at 2000 rpm, washed with saline three times, and then filtered using a 40 μm cell strainer (CLS431750, Corning).

### 5.2 Micro CT imaging

The glaucomatous eyes (8 weeks post-injection) were dissected to remove the lens and collect O-ring-shaped tissue from around the limbus, including iris and cornea. To dehydrate the tissue, it was gradually incubated in ethanol solutions (50% to 100%) for 1 hour with shaking at 120 rpm on an orbital shaker. The dehydrated tissue was then transferred to toluene for 15 minutes, with the solution being replaced with fresh toluene for an additional 15-minute incubation. To improve contrast, the samples were stained with Lugol’s solution in 70% ethanol for 15 minutes, followed by another 15-minute incubation in the same solution. The samples were then rinsed in water for 10 minutes to remove excess contrast reagent. Finally, the tissues were mounted in 2% (w/v) agarose gel in water. X-ray images were obtained using a Zeiss Xradia 620 Versa micro CT, set at 60 kV, 6.5 W, and 108.28 µA. Imaging was conducted with a 20X objective, capturing images every 30 seconds for 15 hours, with a pixel resolution of 0.67 µm and a binning factor of 2 to enhance signal quality. The 3D reconstruction image was generated using Dragonfly 3D world software ver. 2024.1 (Comet Technologies Canada Inc.).

### 5.3 *In vitro* degradation and characterization

To analyze the degradation dynamics of PS/PLGA Viscobeads, Raman spectroscopy was conducted using a Thermo Fisher Scientific DXR2xi Raman microscope equipped with a 780-nm laser and a 10X microscope objective. PS/PLGA Viscobeads were incubated for 7 days in different temperatures at 37°C, and 55°C, respectively. The Viscobeads were then dried on a glass surface. Each spectrum was scanned from 3200 to 500 cm^−1^ with 2 mW laser power and a 25-μm slit width for 5 seconds integration time. Eight scans were done automatically from different spots on the surface and then averaged by the instrument before analysis. To visualize the beads morphology, scanning electron microscopy was operated with dried samples using field-emission scanning electron microscopes (JEOL, JSM-6700F and Hitachi SU5000) under high vacuum conditions at 5 kV and 196-198 µA emission current, maintaining a working distance of 8.16-8.69 mm.

### 5.4 Computational modeling of microbead transport

A two-way coupled computational fluid dynamics (CFD) and discrete element method (DEM) simulation was performed using Ansys Fluent 2023 R2 and Ansys Rocky 2023 R2 to investigate the transport of microbeads through a gradient porous structure. The computational domain consisted of a fluid region partially blocked by three sequential layers of pillar arrays with no-slip boundaries. The pillar diameters decreased from 15 μm to 7 μm to 2 μm along the flow direction, with corresponding inter-pillar distances of 15 μm, 7 μm, and 2 μm, respectively, mimicking the gradient porosity of the TM. The fluid, representing aqueous humor, was modeled as an incompressible, laminar, and transient flow with a density of 998.2 kg/m³ and a viscosity of 1.003 × 10⁻³ kg/(m·s). Fixed-size microbeads with diameters of 2, 4, 6, 8, and 10 μm, as well as heterogeneous microbeads ranging from 1 to 15 μm (Gaussian distribution), were simulated. In all cases, the total mass of injected microbeads was kept constant at 1.83 × 10⁻¹¹ kg. The microbeads were modeled as spherical particles with a density of 1000 kg/m³ and a Young’s modulus of 0.1 GPa using a hysteretic linear spring model. Microbeads were released at the inlet upstream of the porous structure at t = 0, and the simulation was run for 4 ms under constant flow conditions.

To quantify flow obstruction, the pressure drop across the TM-mimicking porous region (p₀–p₃) was calculated. Under constant flow conditions, the pressure drop is proportional to hydraulic resistance (ΔP ∝ R). To relate pore-scale resistance changes to physiologically relevant IOP, a Goldmann-based framework was applied:

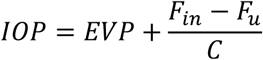

where EVP is episcleral venous pressure, *F_in_* is aqueous humor inflow, *F_u_* is uveoscleral outflow, and *C* is conventional outflow facility. Because *C* ∝ 1/*R*, the relative IOP elevation above EVP can be expressed as proportional to the simulated pressure-drop ratio:

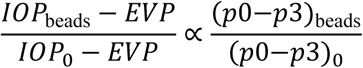

To further estimate absolute IOP values, a calibrated Goldmann-based approximation was used:

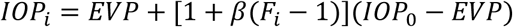

where *F_i_*is the relative pressure-drop factor derived from CFD results, and *β*is a scaling parameter accounting for the partial representation of the outflow pathway. This approach enables a semi-quantitative mapping between bead-induced pore-scale obstruction and expected changes in IOP.

### 5.5 Animals

C57BL/6J wild-type mice, obtained from Jackson Laboratory (JAX), aged between 6 to 8 weeks and weighing 20 to 25 grams were used for this study following the approved animal protocol by the Institutional Animal Care and Use Committee (IACUC) at Binghamton University (Protocol: 894-23) and University of Massachusetts Amherst (Protocol: 2529) and complied with relevant regulations. All mice were housed in cages with a maximum of 5 mice per cage and subjected to a 12 h light/dark cycle, with unrestricted access to food and water.

### 5.6 Intracameral injection of Viscobeads

IOP elevation was induced by injecting Viscobeads into the anterior chamber of both eyes in mice in this study. The surgical procedures were adapted from a well-established microbead occlusion model and our previous published work ^[6a, 13]^. In brief, anesthetized mice had their corneas gently punctured near the center using a 33G needle (CAD4113, Sigma). A bubble was then introduced through this incision site into the anterior chamber to prevent possible leakage. Subsequently, 1 µL of Viscobeads was injected into the anterior chamber. After a 5- minute interval during which the Viscobeads accumulated at the iridocorneal angle, the mice were treated with antibiotic Vetropolycin ointment (Dechra Veterinary Products) and placed on a heating pad for recovery.

### 5.7 Design of overexpression and knockdown vectors

Following previously established protocols ^[13, 33]^, viral vectors were designed and synthesized by Synbio Technologies (Monmouth Junction, NJ) using the pAAV-hSyn-hChR2(H134R)-EYFP plasmid (Addgene #26973) as a backbone. The hChR2(H134R)-EYFP sequence was replaced with the target gene sequences of DCLK2, ARMCX1, LIN28, or IGF-1 for overexpression experiments. AAV2/2 viral particles were subsequently produced by the Boston Children’s Hospital Viral Core. For CRISPR-mediated gene knockout, a modified AAV-U6-sgRNA-hSyn-mCherry plasmid (Addgene #87916) was used. To minimize off-target effects, we delivered a combination of five AAV2 single-guide RNA (sgRNA) expression vectors along with AAV2-Cas9 into the eyes of wild-type mice, achieving high infection efficiency as previously described ^[33–34]^. The vectors and sequences used in these manipulation experiments are listed in **Table 1**.

**Table 1:**
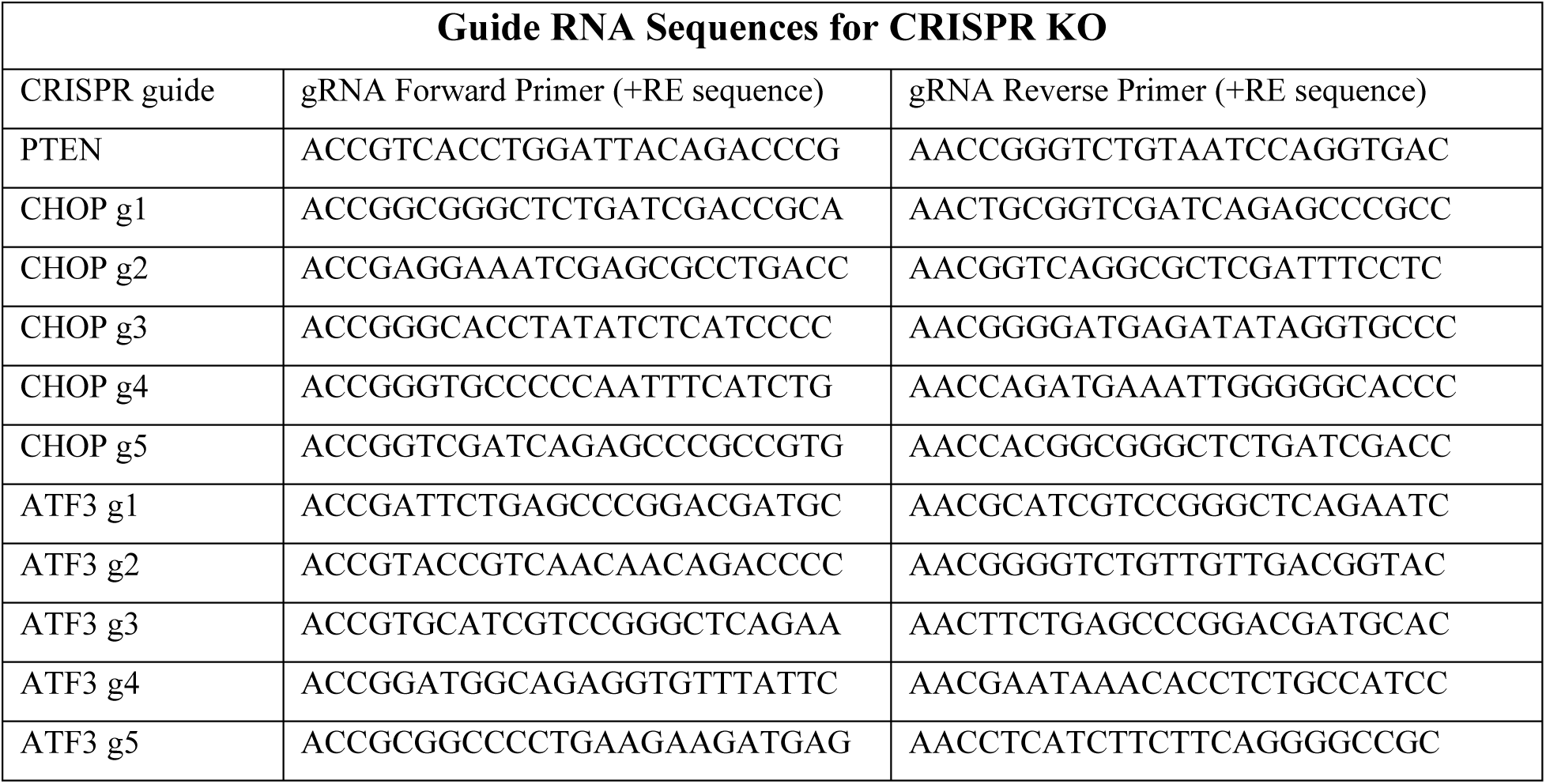
Guide RNA Sequences used in our work for CRISPR knockout (KO)

### 5.8 Intravitreal injection of AAV

AAV vectors containing various gene candidates were delivered via intravitreal injection into mice eyes. A total 2 µL of the AAV virus was injected into the vitreous with each AAV titer adjusted to 1 ✕ 10^13^ GC/mL. Specifically, 2 µL of AAV2/2-hsyn-DCLK2, ARMCX1 or LIN28 were injected in C57BL/6J mice to overexpress genes. For the LIN28 and IGF-1 combination, AAV2/2-hsyn-LIN28 and IGF-1 were mixed at a 1:1 ratio. For CRISPR knockout targeting each gene candidate (PTEN, ATF3, CHOP, non-target), AAV2/2-sgRNA and Cas9 were also mixed at a 1:1 ratio. Briefly, a pulled-glass micropipette was inserted near the peripheral retina behind the ora serrata and deliberately angled to avoid damage to the lens. 2 µL of the AAV virus were injected into mice vitreous. Following the injection, we applied antibiotic ophthalmic ointment to the corneal surface.

### 5.9 Intraocular pressure measurement

IOP measurements were performed using a TonoLab tonometer (Colonial Medical Supply) according to the manufacturer’s instructions. Because chronic IOP measurements in awake mice are technically challenging due to eye movements, stress responses, and variability in probe alignment, all measurements were performed under a standardized isoflurane anesthesia protocol (3% isoflurane in air). Although anesthesia can reduce absolute IOP values, the same anesthetic conditions were applied consistently across all experimental groups to ensure comparability of relative differences. To minimize the acute effect of isoflurane on IOP, measurements were taken 5 minutes after loss of the lid reflex to allow partial pressure stabilization. The average IOP was calculated from five consecutive readings, excluding the highest and lowest values. All measurements were performed in a blinded manner.

### 5.10 Pattern electroretinogram (pERG) recording

Pattern ERG recording was performed on mice before surgery, 2, 4, and 6 weeks after surgery using the Celeris Diagnosys system (Diagnosys LLC). Anesthesia was administered, and the mice were positioned on a heating table to maintain body temperature. Pupillary dilation was achieved using 1% tropicamide. Room illumination was maintained with dim red light. Gonak gel (Akorn, Inc.) was applied to the cornea of each eye before electrode placement. ERG was recorded from the corneal surface using Diagnosys Celeris pattern stimulator electrodes while simultaneously presenting visual stimulation. Reference and ground electrodes are small stainless-steel needles inserted in the skin of the forehead area and the base of the tail, respectively. Prior to recording, electrode impedance was measured, ensuring electrodes had impedance values between 5-7 kΩ.

Horizontal contrast-reversing gratings were presented via Diagnosys Celeris pattern stimulator electrodes, with a visual stimulus contrast of 95%, luminance of 50 cd m^-^², spatial resolution of 0.052 cycles/degree, and a reversal frequency of 2.1 Hz. Each data trace captured lasted for 470 ms, including a 50 ms baseline prior to stimulus onset. The signal was sampled at a frequency of 2000 Hz. The average of two consecutive recordings of 300 traces each was taken to produce one reading. The pERG measurements included the mean amplitudes of the P1 (first positive peak) and N2 (second negative peak) components. The P1 component represents the initial positive deflection, originating from RGCs and outer retinal photoreceptors. The mean P1 amplitude was measured as the difference between the average lowest point of the N1 component and the average peak of the P1 component. The N2 component is the negative deflection following the P1 component, originating from the inner retina and reflecting RGC function. The mean N2 amplitude was calculated as the difference between the average peak of the P1 component and the average lowest point of the N2 component.

### 5.11 Flash visual evoked potential (VEP)

Flash VEP testing was performed on mice prior to surgery and at 2, 4, and 6 weeks post-Viscobead injection using the Celeris Diagnosys system (Diagnosys LLC.). Mice were dark-adapted overnight before the VEP recordings. Anesthesia was administered, and the mice were positioned on a heating table to maintain body temperature. Pupillary dilation was achieved using 1% tropicamide. Room illumination was maintained with dim red light. Gonak gel (Akorn, Inc.) was applied to the cornea of each eye before Diagnosys Celeris touch stimulator placement. A reference needle electrode for VEP was placed in the cheek, while the active VEP electrode was positioned at the midline on the back of the head, close to the location of the visual cortex. The ground electrode was placed in the hindquarters near the tail. We adhered to the standard protocols of the International Society for Clinical Electrophysiology of Vision (ISCEV) ^[35]^. Prior to recording, electrode impedance was measured, ensuring recording electrodes had impedance values below 2 kΩ. White flash stimulation was delivered using the Diagnosys Celeris touch stimulator electrodes at an intensity of 1 cd s^-1^ m^-2^. Each stimulus lasted for 5 ms with an interval of 10 s. Each recorded trace had a duration of 320 ms, including a 20 ms pre-stimulus baseline. The signals were amplified 1,000 times with a bandpass filter set between 0.1 Hz and 300 Hz. Each waveform was averaged over 61 superimposed responses to enhance signal-to-noise ratio. Flash VEP inspection was performed using a two-channel recording method and measured at least three times consecutively. The flash VEP measurement included the mean amplitude and implicit time of the mean responsive curve. The VEP amplitude was defined as the difference between this baseline and the first negative peak within the first 300 ms following the onset of the stimulus. The implicit time was determined as the interval from the onset of the light stimulus to the point when the first negative peak was reached.

### 5.12 Immunohistochemistry

For immunostaining, the animals were euthanized with an overdose of anesthesia, and perfused transcardially with ice-cold phosphate buffer saline (PBS, P3813, Sigma), followed by 4% paraformaldehyde (PFA, P6148, Sigma). Subsequently, the optic nerves were carefully dissected and postfixed in 4% PFA overnight at 4°C. For the retinal wholemount, the retinal tissues were carefully extracted from the eyecup and flattened with four flower-shaped radial incisions. To ensure cryoprotection, the tissues were immersed in a 30% sucrose solution in PBS for 48 hours. After being embedded in the Optimal Cutting Temperature compound (Tissue Tek, Sakura Finetek) with dry ice, the samples were cut into 12 μm sections for the optic nerves and 20 μm for retina tissues. Cryosectioned samples and wholemount retinas were rinsed in PBS and subsequently blocked in a solution containing PBS, 1% Triton X- 100 (X100, Sigma), and 5% horse serum (Gibco, as whole-mount buffer) overnight at 4°C. Primary antibodies, diluted in the whole-mount buffer, were applied for 2-4 days at 4°C, followed by three washes with PBS (each lasting 10 minutes). Secondary antibodies, all diluted at a 1:1000 ratio in PBS, were applied and left overnight at 4°C. After five washes with PBS (each lasting 10 minutes), the retinas were mounted using Fluoromount-G (0100- 01, Southern Biotech). Confocal images were collected with Leica SP2, Zeiss 700 or Zeiss 710 confocal microscopes. Image stacking and quantitative analysis of RGC loss were processed by Fiji software or CellCount software. Specifically, we examined the entire cross-sectioned retina tissues to count RGC actress three different slices per eye using a Nikon inverted microscope. We then extracted images to label the RBPMS channel for individual RGC identification, and used Cell Counter software to automatically count the cells and calculate the average number.

### 5.13 Sleep pattern monitoring

Mice were housed in the PiezoSleep™ system (Signal solutions, LLC.), which has shown 92% accuracy in classifying asleep and awake states compared to the traditional EEG/EMG method ^[25–26]^. The mice were continuously monitored for 24 hours to capture their sleep patterns, with housing cages featuring automated lights settings (a 12-hour light/dark cycle). The recorded data were analyzed using a custom MATLAB algorithm to reduce noise and artifacts, eliminate breath rate interference, and identify different sleeping states (awake, NREM, REM).

### 5.14 Data analysis

Normality and variance similarity were assessed using Microsoft Excel and the R programming language before applying any parametric tests. If the criteria for parametric tests were not met, non-parametric tests were carried out. For single comparisons between two groups, a two-tailed Student’s t-test was utilized. In cases of multiple groups, the data were analyzed using either one-way or two-way ANOVA, depending on the appropriate experimental design. Post hoc comparisons were performed only when statistical significance was observed in the primary measure, and Bonferroni’s correction was used to adjust the p-values for multiple comparisons. Error bars in all figures represent the mean ± S.E.M. The mice, which had varying litters, body weights, and sexes, were randomized and assigned to different treatment groups. No additional randomization methods were employed for the animal studies. Each experimental data value represents individual measurements or observations, and their descriptions are provided in the corresponding figure captions. All statistical analyses were conducted using GraphPad Prism 10. Significance was determined with P values less than 0.05, denoted as follows: *0.01≤p<0.05, **0.001≤p<0.01, ***0.0001≤p<0.001, ****p<0.0001, and “ns” for not significant.

## Author contributions

Z.H., S.R., and QB.W. conceptualized the Viscobeads idea and designed all the experiments. E.H., S.H., and QB.W. took the lead in fabricating and characterizing the Viscobeads *in vitro* and *in vivo*. F.T. and S.Z. contributed to the gene therapy part. QL.W. and R.Y. contributed to the recordings. E.H., QL.W., R.D., S.T., C.L., M.X and QB.W. managed cellular biocompatibility analysis, glaucoma surgeries, IOP measurements, and immunohistochemistry. E.H. and L.Y. characterized the trabecular meshwork. X.L., J.T., and Y.W. assisted with the simulation. S.T., R.D. and E.H. assisted with tissue harvesting. E.H., R.Y., S.R. and QB.W. wrote the manuscript with input from all authors.

## Acknowledgments

This study was supported by the National Institutes of Health (R00MH120279, SR; R21NS144777), the Air Force Office of Scientific Research Young Investigator Program (FA9550-23-1-0339, SR), the National Science Foundation Faculty Early Career Development Program (CAREER; 2414753, SR; 2442112, QBW), the Craig H. Neilsen Foundation SCIRTS Pilot Research Grant (1348565), the Brain & Behavior Research Foundation Young Investigator Grant (29878, SR), the S.H. Ho Foundation Research Grants for Health Sciences and Technology, Faculty Startup Funds from Binghamton University, Binghamton University Transdisciplinary Areas of Excellence (TAE) program (QBW), the Binghamton University Watson College of Engineering and Applied Science Seed Grant Program (105383), and the Binghamton University S3IP Small Grant Awards (ADLG 267 and ADLG 324, SR; ADLG274, QBW). Additionally, we acknowledge the crucial resources offered by both the UMass Amherst and Binghamton core facilities and Animal Care Service departments.

## Data Availability Statement

All data are available in the main text or Supporting Information, with additional data available upon request.

