## supplementary information for "Biologically Driven *In Vivo* Occlusion Design Provides a Reliable Experimental Glaucoma Mouse Model"

**Supplementary Methods**

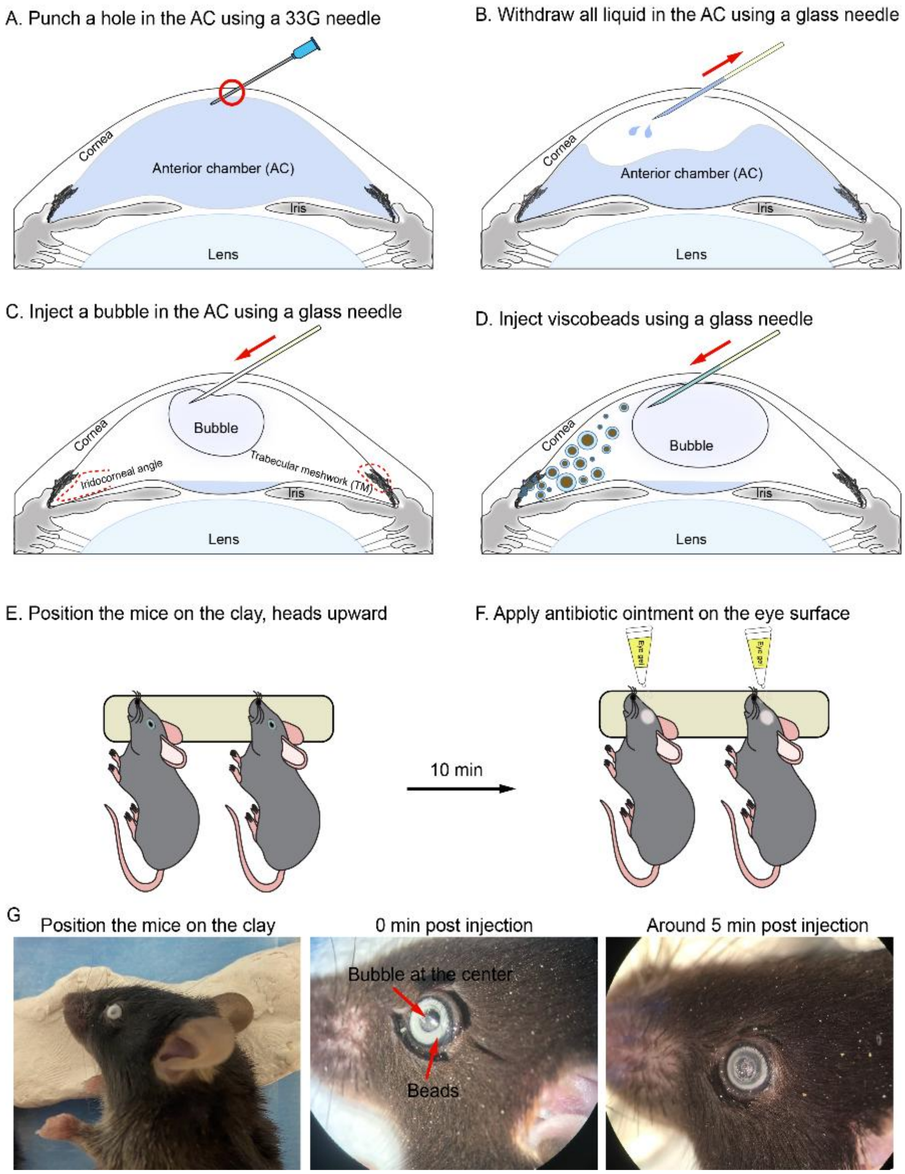


**Figure S1**: Intracameral injection procedure: (A) Create a central corneal hole using a 33G needle to minimize leakage. (B) Insert a glass microneedle at ~45° to the cornea, remove aqueous humor, and separate the iris from the cornea. (C) Inject a large air bubble to keep the iris and cornea apart, ensuring it remains in the anterior chamber. (D) Inject 1 µL of vortexed microbead solution (33% v/v in saline or PBS) using a glass microneedle (tip OD 70–120 µm). Maintain an intact air bubble to prevent leakage. (E) Position the mouse with the injected eye facing upward at water level to ensure uniform microbead distribution in the trabecular meshwork. (F) After 10 minutes, apply antibiotic eye gel. If both eyes require injection, treat the second eye on the following day. Mice recover on a heated plate. (G) Representative photographs of a mouse placed on the clay with the injected eye facing upward at water level (left), the Viscobeads injected into the anterior chamber (middle), and the Viscobeads accumulated at the mouse's iridocorneal angle 5 minutes after injection (right). It is noted that the air bubble remained intact in the central anterior chamber to prevent microbead leakage.

### 4. Biocompatibility tests on mouse corneal cells

Primary mouse corneal epithelial cells were isolated from postnatal day 7 (P7) or earlier C57BL/6 mouse corneas and cultured in Matrigel-coated 48-well plates using corneal epithelial cell medium (ATCC, catalog no. PCS-700-040 and PCS-700-030). TM cells were isolated from P7 or earlier C57BL/6 mice and cultured in Matrigel-coated 24-well plates using TM cell growth medium consisting of MEM supplemented with 1% GlutaMAX™, 10% FBS, and 1% penicillin-streptomycin. Neuro-2A cells were purchased from ATCC (catalog no. CCL-131) and cultured in Dulbecco's Modified Eagle Medium (DMEM) supplemented with 10% fetal bovine serum (FBS) and 1% penicillin-streptomycin. Cell viability was evaluated using Calcein-AM (2 µL of 1 mg/mL per well; green fluorescence, Sigma-Aldrich, catalog no. 17783) to label live cells. Dead cells were stained with Ethidium Homodimer-1 (2 µL of 1 mg/mL per well; red fluorescence, Sigma-Aldrich, catalog no. 46043). Fluorescent images were acquired using an inverted fluorescence microscope (Eclipse, Nikon, Melville, NY) to capture signals from cells under different treatment conditions. Quantitative analysis of live and dead cells was performed using ImageJ software. Cell viability was calculated as the ratio of live cells to total cells (live + dead) using the following formula:

$$Cell\ viability\ (\%) = \frac{Viable\ cell\ number}{Total\ cell\ numbers} \times 100\%$$

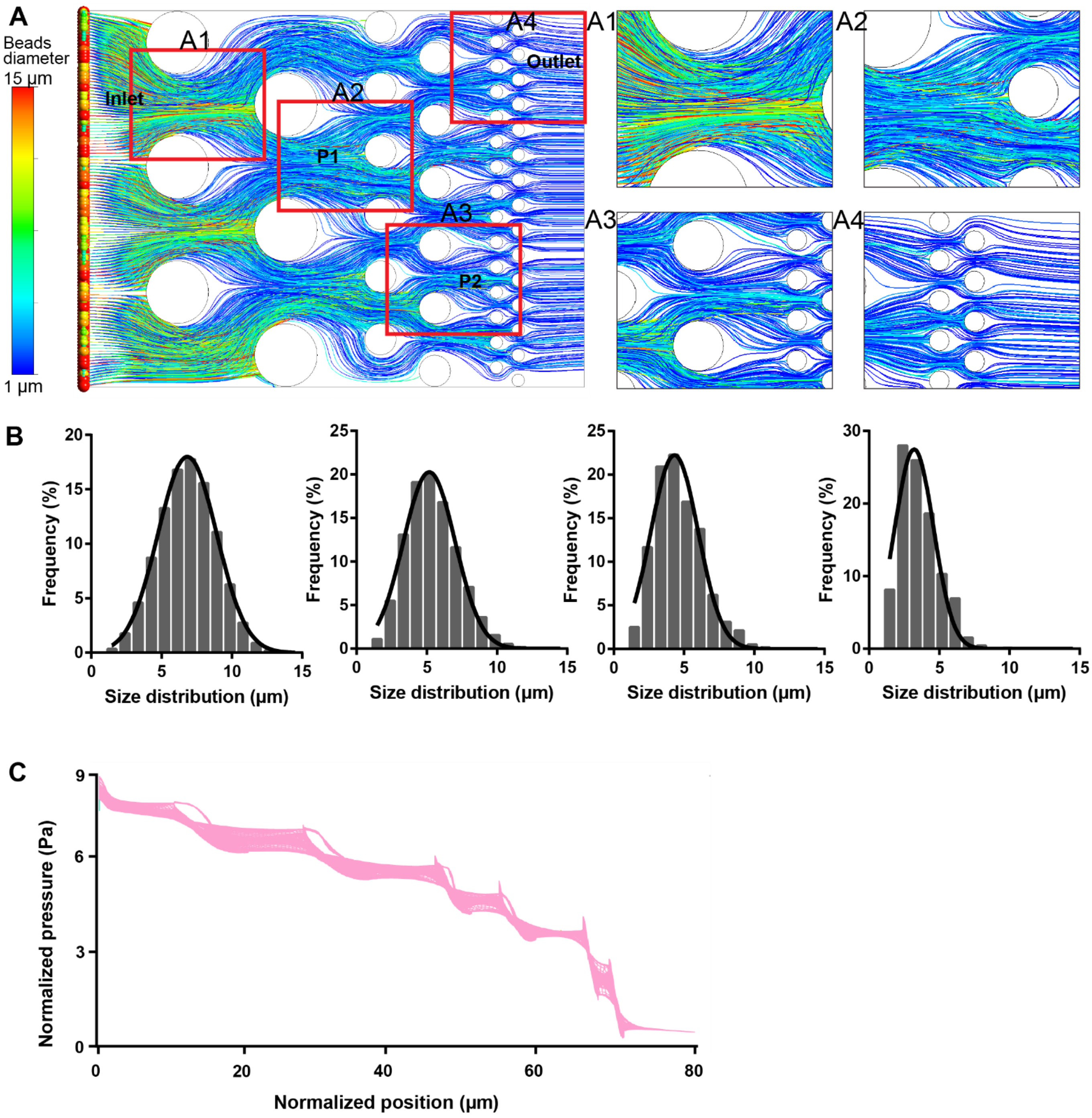


**Fig. S2:** (A) The simulation of physical blockage in aqueous humor outflow was conducted using ANSYS Fluent. Heterogeneously distributed particles (Gaussian distribution with sizes ranging from 1 to 15 µm) were released from the inlet of a three-layered structure with gradient gap sizes (10, 5, and 2 µm from inlet to outlet, respectively) in a discrete phase model. The color lines indicate trajectories of particles with different sizes. (A1-A4) Enlarged views show the outflow of the fluid with particles sized 1-15 µm at different layers (inlet: A1, P1: A2, P2: A3, and outlet: A4). (B) The size distribution of particles at different layers (inlet, P1, P2, and outlet) in the ANSYS Fluent model indicates particle entrapment in distinct layers. (C) Normalized pressure changes at different positions from the inlet to the outlet.

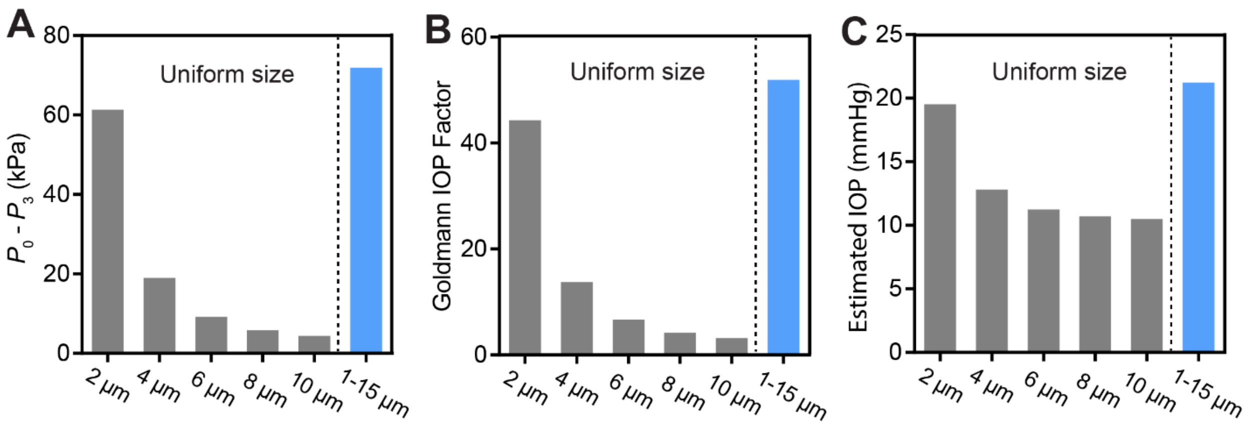


**Fig. S3:** Goldmann-based analysis linking CFD-simulated local obstruction to predicted IOP changes. (a) Simulated pressure drop across the TM-mimicking porous structure (p0–p3) over time for the 2 µm, 4 µm, 6 µm, 8 µm, and 10 µm uniform beads condition and the heterogeneous bead population (1–15 µm). (b) Goldmann-style relative IOP factors calculated from the p0-p3-based pressure-drop ratios. (c) Estimated absolute IOP values obtained using a calibrated Goldmann-based model ($IOP_0 = 10$ mmHg, $EVP = 8$mmHg, $\beta = 0.11$).

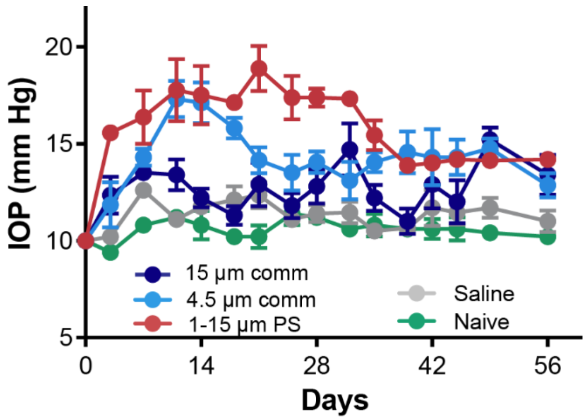


**Fig. S4:** Intraocular pressure in mice injected with 1-15 µm PS microbeads, 15 µm and 4.5 µm commercialized (comm) PS microbeads, saline, and the naïve group. n= 4 eyes for 1-15 µm PS, 5 eyes for other groups.

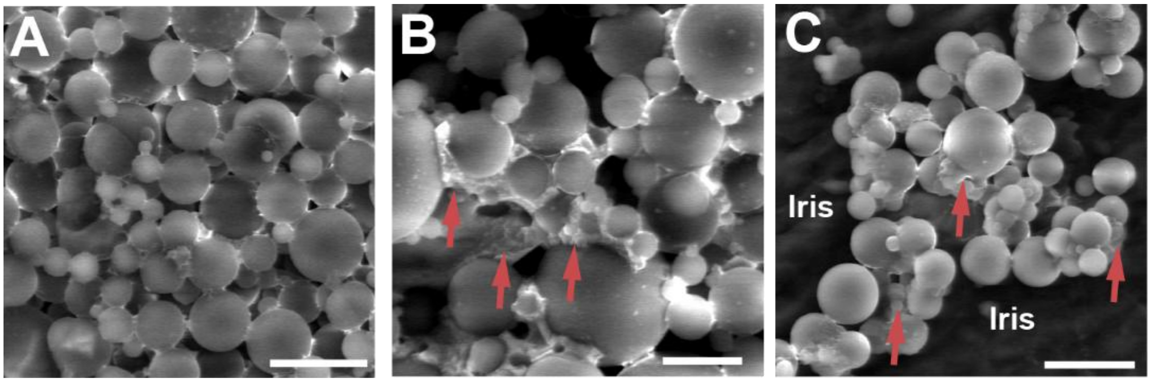


**Fig. S5:** Representative SEM images of PS/PLGA Viscobeads before injection (A) and after 8 weeks of injection (B-C) suggest that degraded PLGA could function as a viscous substance, efficiently adhering individual microbeads in vivo. This is confirmed by SEM observations of the remaining beads on the iris surface.

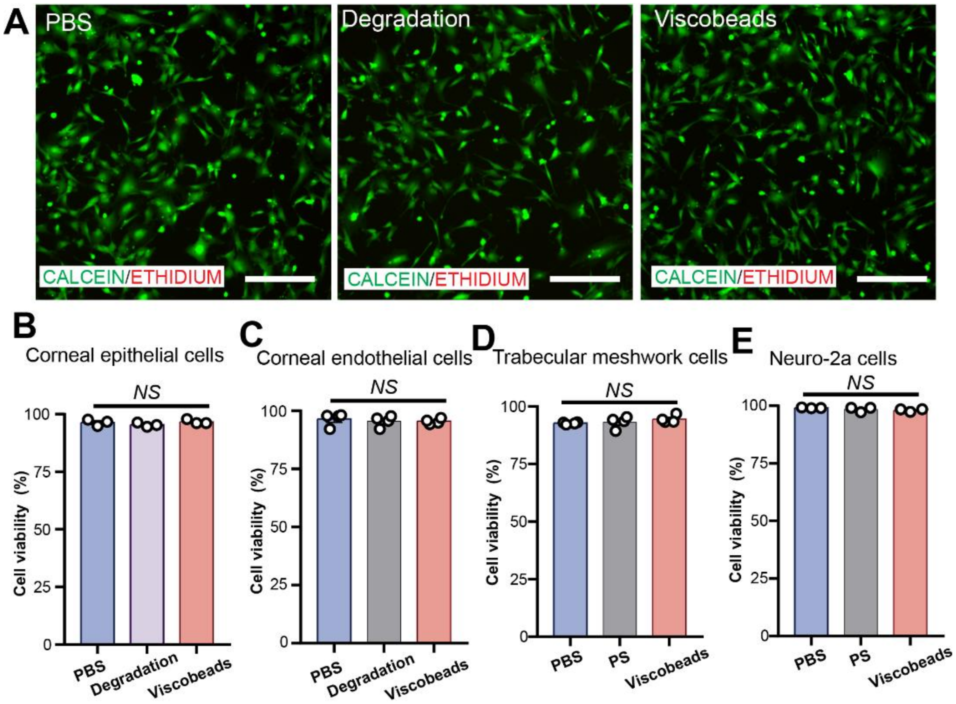


**Fig. S6:** Biocompatibility of PS/PLGA viscobeads and their degradation byproducts in different cell lines. (A) Representative fluorescence microscopy images of mouse corneal epithelial cells following treatment with PS/PLGA viscobeads or their degradation byproducts. Live cells are stained green with Calcein-AM, and dead cells are stained red with ethidium homodimer-1. (B) Quantification of corneal epithelial cell viability (n=3 independent samples). (C) Quantification of corneal endothelial cells (n=4 independent samples) (D) Quantification of trabecular meshwork cell viability (n=4 independent samples). (E) Quantification of Neuro-2A cell viability. Data are shown as mean ± SEM (n=3 independent samples). NS, not significant. Scale bar, 200 μm.

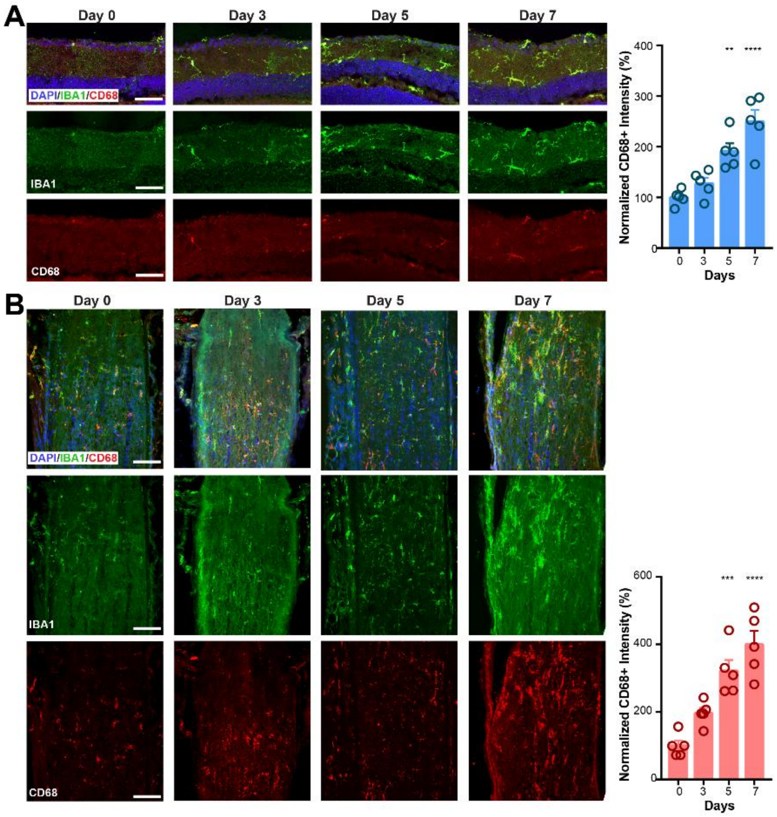


**Fig. S7:** Representative confocal images (left) and quantification (right) of retinal cryosections (A) and optic nerve head (B) stained with DAPI, IBA1, and CD68 at days 0, 3, 5, and 7 post-injection. Scale bars: 50 μm.

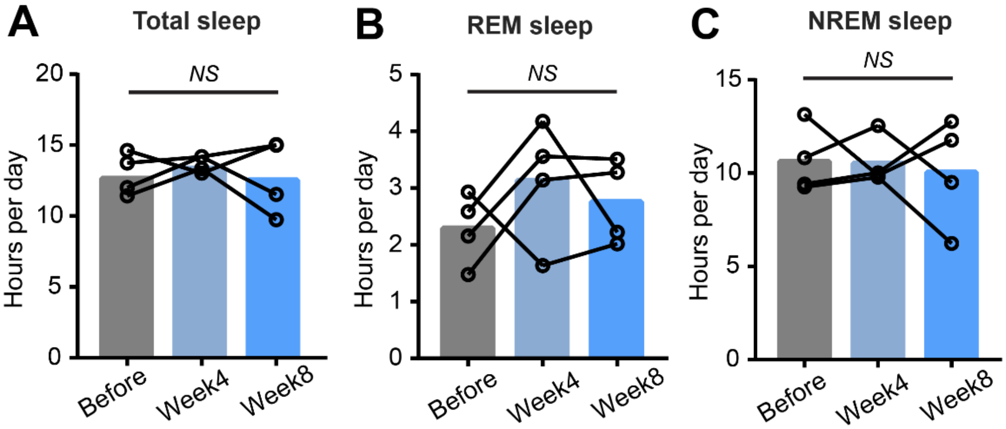


**Fig. S8:** Total sleep (A), REM sleep (B), and NREM sleep (C) duration per 24 hours in mice before saline injection, and at 4 and 8 weeks post-injection of saline. n=4 mice.
